# Studies on the *Escherichia coli* ExbD Transmembrane Domain, Residue L132, and an Inhibitory Cyclic Peptide

**DOI:** 10.1101/2022.09.26.509584

**Authors:** Bimal Jana, Dale Kopp, Mingchao Xie, Hema Vakharia-Rao, Kathleen Postle

## Abstract

The TonB system of *Escherichia coli* uses the cytoplasmic membrane protonmotive force (PMF) to energize active transport of nutrients across the otherwise unenergized outer membrane. Because it overcomes limitations for nutrient diffusion through outer membrane size-limiting porins, it provides a growth advantage and is widespread among Gram-negative bacteria. It consists of three known cytoplasmic membrane proteins, TonB, ExbB and ExbD that energize a variety of customized TonB-dependent transporters in the outer membrane. The sole ExbD transmembrane domain is proposed to consist of residues 23-43 (Kampfenkel and Braun, 1992, J. Bacteriol. 174:5485-7). Here we showed that the charge and location of residue Asp25 were essential for activity of the TonB system, thus identifying it as the only PMF-responsive element in the TonB system. The proposed boundaries of the transmembrane domain α-helix were revised to consist of residues 23-39, with residues 40-43 initiating the subsequent disordered region required for signal transduction (Kopp and Postle, 2020, J. Bacteriol. 202, e00687-19). Trapping of disulfide-linked ExbD homodimers through T42C or V43C prevented TonB system activity that was restored by addition of the reducing agent dithiothreitol, indicating a requirement for motion. In *vivo* photo-cross-linking experiments suggested that motion was rotation of ExbD transmembrane domains. Inactivity of ExbD L132Q, the first ExbD mutant identified, was likely due to steric hindrance. A conserved and defined site of *in vivo* ExbD interaction with TonB was identified. Exogenous addition of a cyclic peptide based on that site inhibited ExbD-TonB interaction while concomitantly decreasing iron transport efficiency. This suggested that a novel antimicrobial strategy against ESKAPE and other Gram-negative pathogens could be developed by targeting ExbD protein-protein interactions.

## INTRODUCTION

Gram-negative bacteria are characterized by their distinctive outer membranes that serve as intrinsic barriers to environmental agents, antibiotics, and degradative enzymes. To overcome the concomitant nutrient limitations imposed by the outer membrane, the TonB system energizes transport across the outer membrane of a wide array of scarce, important nutrients such as iron-siderophores, heme, zinc, copper, cobalamin, nickel, maltodextrin, chitin oligosaccharides, and sucrose [for reviews see (1–8)]. The TonB system is also an essential component for pathogenesis by certain Gram-negative bacteria (9).

In *Escherichia coli* K12 the TonB-dependent transport ligands are iron-siderophores and cobalamin. The TonB system consists of integral cytoplasmic membrane proteins ExbB and ExbD, which harvest the protonmotive force (PMF) of the cytoplasmic membrane, and transmit it to the integral cytoplasmic membrane protein TonB, which then transmits mechanical energy to customized β-barrels in the outer membrane known as TonB-dependent transporters (TBDTs) (Fig.1). In the absence of PMF or TonB, ligands bind to TBDTs but do not cross the outer membrane or enter the periplasm. Because the TBDTs greatly outnumber TonB and compete for contact with it, it is likely that TonB engages in an energy transduction cycle involving transient interactions with individual TBDTs (10–12).

**Figure 1.**
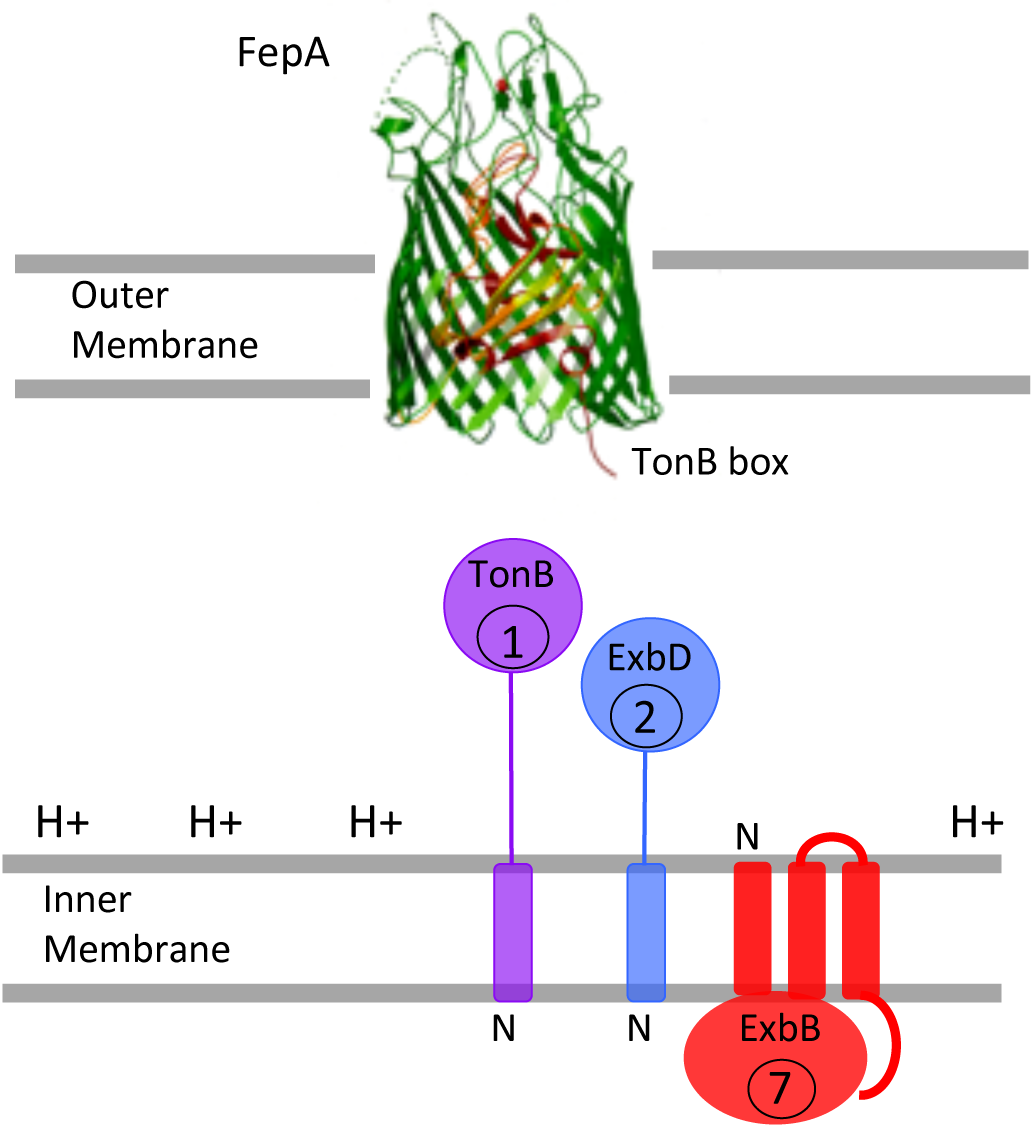
The TonB system of *Escherichia coli* K12. The TonB-dependent transporter, FepA, is shown in the outer membrane. At its extreme amino terminus, the TonB box, a site of interaction with TonB, is shown protruding into the periplasm. The topologies of the cytoplasmic membrane proteins TonB, ExbB and ExbD are shown in the cytoplasmic membrane. Cellular ratios of the three proteins are indicated in circles in each protein. The protonmotive force gradient of the cytoplasmic membrane (H+) is shown. The crystal structure of FepA was solved by Buchanan et al. (120).

Structurally, the TBDT β-barrels consist of 22 strands, with a lumen that is occluded by an amino-terminal globular domain, called the cork or hatch. *In vivo*, the periplasmically exposed surface of the TBDT appears to signal that ligand is bound via a conformational change that allows for productive interaction with TonB (13–18). The cork domain moves *in vivo* during contact with TonB (18).

Among the three integral cytoplasmic membrane proteins in the TonB system, there are five transmembrane domains, any one of which could theoretically respond to the PMF of the cytoplasmic membrane. TonB and ExbD are both anchored by a single transmembrane domain with the majority of each protein occupying the periplasmic space (Fig.1) [(19–21). ExbB has three transmembrane domains, with an N-out, C-in topology and primarily occupies the cytoplasm (22). TonB and ExbD both depend on ExbB for their proteolytic stability suggesting that ExbB constitutes a scaffolding for their assembly [(23–25); unpublished results]. Consistent with that idea, the cellular ratio of TonB:ExbD:ExbB is 1:2:7 (11). The TonB transmembrane domain remains homodimerized throughout the energy transduction cycle, thus making the ratio 2:4:14 (12). Our current model suggests that, as TonB initiates an energy transduction cycle, its previously monomeric periplasmic domain becomes a homodimer, that can then interact with ExbD homodimerized through its periplasmic domain; this transforms into a TonB-ExbD heterodimer that subsequently configures TonB to energize active transport of nutrients through a TBDT (Fig. 2). In this model the proposed energy-dependent step is recycling of ExbD homodimers to an energized state.

**Figure 2.**
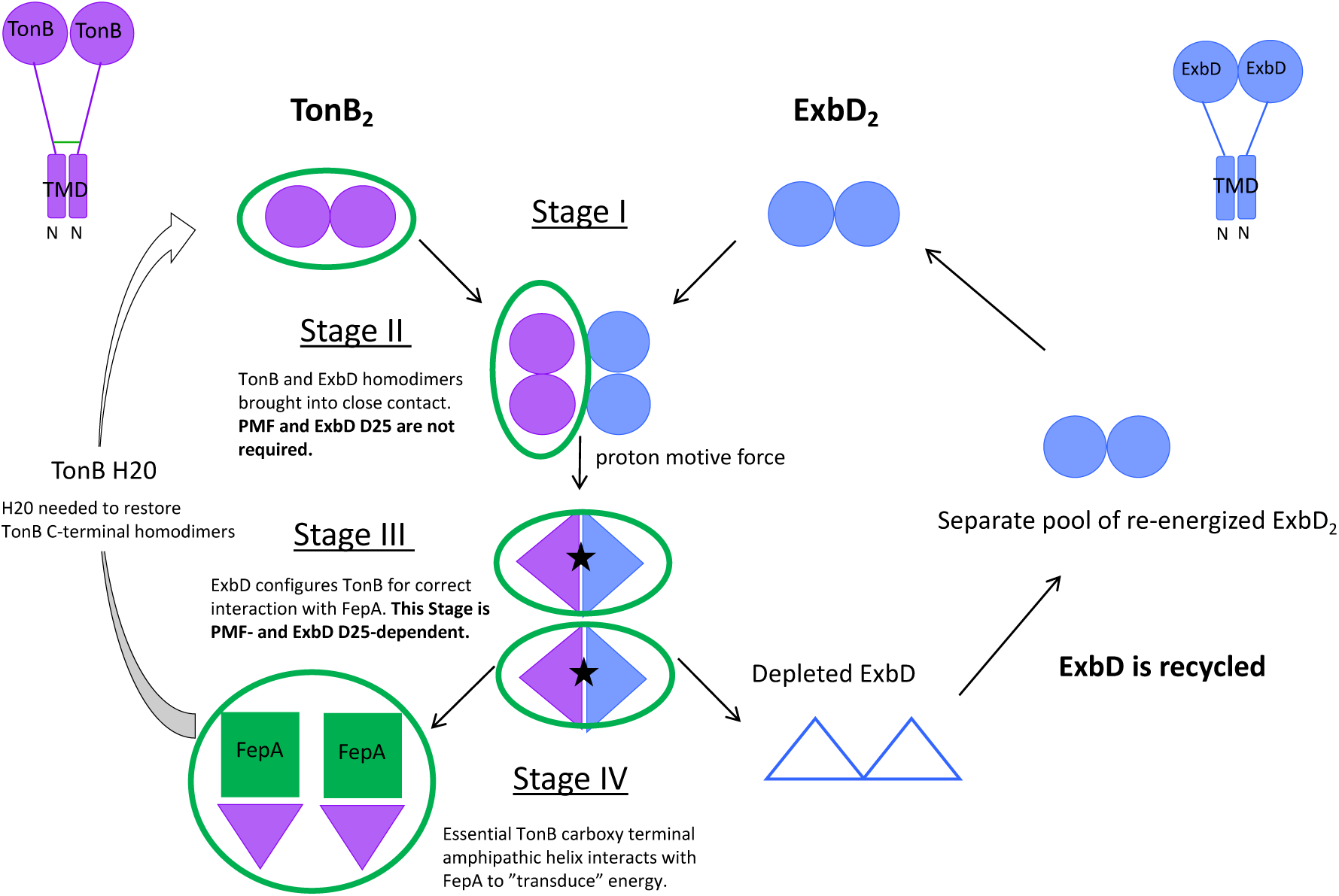
Model for the TonB energy transduction cycle and the role of ExbD based on *in vivo* data [adapted from (12)]. Full length TonB (purple, upper left corner) and ExbD (blue, upper right corner) each have a single transmembrane domain signal anchor in the cytoplasmic membrane with the bulk of the residues occupying the periplasmic space. The TonB transmembrane domain remains homodimerized throughout the cycle (12). The ExbD transmembrane domain also homodimerizes (as shown in this work), but it is not known if this persists throughout the cycle. Interactions between the periplasmic carboxy-terminal domains of TonB_2_ and ExbD_2_ homodimers are shown. Purple circles/triangles are TonB carboxy termini. Blue circles/triangles are ExbD carboxy termini. ExbB tetramers (ExbB_4_), shown to exist *in vivo*, independently stabilize both TonB and ExbD (23, 80). For simplicity, ExbB_4_ is not shown in this model. The TonB-dependent transporter, FepA, is represented by the green square. In Stage I, TonB periplasmic domains form obligatory homodimers through residues in and near its single carboxy terminal amphipathic helix (residues 199-216), mediated by H20 in the TonB TMD, which plays a structural role (12, 26, 41, 72). ExbD periplasmic domains also homodimerize (36). In Stage I there is no evidence that TonB and ExbD homodimers interact, yet both are proteolytically stabilized by ExbB, suggesting that they might exist in separate ExbB tetramers. Such a distribution also helps to account for cellular ratios of 14 ExbB: 4ExbD: 2TonB. In Stage II TonB and ExbD periplasmic domain homodimers are brought into close contact by movement of the ExbB tetramers such that ExbD protects TonB in spheroplasts from degradation by exogenously added protease (36, 42). In Stage III, the previously homodimeric TonBs and ExbD periplasmic domains convert into a dimer of TonB-ExbD heterodimers in spheroplasts that become sensitive to degradation by exogenously added protease. Both PMF and ExbD D25 are required (36, 42). Energized ExbD correctly configures TonB for productive interaction with FepA. Stage IV is binding of a monomeric periplasmic domain of TonB to FepA, such that the siderophore enterochelin is actively transported across the outer membrane (18). After a transport event, H20, a key structural residue in the TonB transmembrane domain, is somehow required for re-establishment of TonB periplasmic domain homodimers in Stage I (12, 26, 41). The green circles represent that all stages of the TonB energy transduction cycle involve the participation of the essential carboxy terminal amphipathic helix (residues 199-216) in TonB (18). In this model, the cellular ratios of 14 ExbB: 4ExbD: 2TonB leads to an excess of 2 ExbDs that are not associated with TonB (11). Because TonB functions as an obligate homodimer through its transmembrane domain (12), we speculate that ExbD_2_ is “de-energized” after Stage IV (empty blue triangles) and is separately recycled. In this model, ExbD_2_ moves in and out of interactions with TonB_2_ escorted by ExbB_4_ tetramers. A separate pool of recycled ExbB_4_-ExbD_2_ is hypothesized to exist to replenish Stage I ExbB_4_-ExbD_2_ homodimers. See (12) for a full explanation of the experimental basis for the model.

The sole potentially PMF-responsive residue in the TonB transmembrane domain, His20, can be fully functionally replaced by a non-protonatable Asparagine residue, thus removing it from consideration as part of the mechanism of energy transduction (26). Likewise, all of the potentially PMF-responsive residues in the three ExbB transmembrane domains are excluded because non-protonatable substitutions are active (27). This leaves Asp25 in the ExbD transmembrane domain as the sole actor that could potentially mediate PMF-dependence in the TonB system.

Because both active and inactive TonBs can bind to TBDTs that are both ligand-bound and ligand-free both *in vivo* and *in vitro*, energy transduction cannot be explained by simple binding (14, 28–34). Only *in vivo* full-length active TonB leads to active transport of ligands through a TBDT, suggesting that there must be a difference in the residues through which the active and inactive TonBs bind TBDTs. That difference appears to be manifested by the ability of TonB to interact correctly with PMF-activated ExbD such that it achieves an “energized” conformation.

10-residue deletion scanning suggests that all ExbD residues are functionally important except for a few amino acids at the cytoplasmic amino terminus (35). ExbD makes multiple *in vivo* contacts during TonB-dependent energy transduction, including formaldehyde cross-linked ExbD-ExbD homodimers, and ExbD-ExbB and ExbD-TonB heterodimers. Consistent with the formaldehyde cross-linking studies, the ExbD periplasmic domain forms a disulfide cross-linked ExbD-ExbD homodimer and ExbD-TonB heterodimer *in vivo*, with nearly the same ExbD residues involved in both complexes (36). In contrast, TonB is essentially impervious to missense mutations, with H20 in the transmembrane domain playing only a structural role [(26, 37–41); Potter and Postle, unpublished results]. Thus, TonB appears to be a puppet with ExbD as the puppet-master.

The role of ExbD can be recapitulated in spheroplasts, which retain their PMF, as well as all of the elements of the cytoplasm that the TonB system may require. The conformation of ExbD is PMF-dependent and it is only by interaction with ExbD that TonB conformations respond to PMF (42).

There have been several indications as to the importance of ExbD Asp 25. ExbD D25N inactivates ExbD (and thus the entire TonB system) and is dominant negative, consistent with being part of a multiprotein complex (43). ExbD D25N mimics the effect of PMF collapse that prevents the ExbD-TonB interaction characterized by *in vivo* formaldehyde cross-linking (29). In the proposed TonB energy transduction cycle, ExbD D25N stalls the ExbD-TonB periplasmic domain interaction at Stage II, the energy-<u>in</u>dependent stage, such that it cannot proceed to the energy-dependent ExbD-TonB interaction [Fig. 2; (29, 36, 42–44)]. Like wild-type ExbD, ExbD D25N formaldehyde-cross-links normally with ExbB monomer and forms homodimers *in vivo* (29).

Along with ExbD, MotB and TolQ participate in three paralogous cytoplasmic membrane systems in *E. coli* K12. Each of the systems uses PMF to energize events at the outer membrane and has two cytoplasmic membrane components in common: one protein with multiple transmembrane domains, occupying primarily the cytoplasm (ExbB, TolQ, and MotA), and one protein anchored by a single transmembrane domain, occupying primarily the periplasm (ExbD, TolR, and MotB) (45–48). TolR and MotB carry an essential Asp residue in their single transmembrane domain at positions 23 and 32 respectively (47, 49), suggesting that ExbD Asp 25 might be equally important.

Here we present a detailed analysis of the ExbD transmembrane domain indicating the pivotal role both charge and location of Asp25 play in TonB system response to cytoplasmic membrane PMF. We redefine the boundaries of the transmembrane domain and identify an subsequent adjacent segment that initiates a previously characterized disordered domain. We show that the ExbD transmembrane domains homodimerize, appear to rotate relative to one another, and that, if dynamic movement is prevented by disulfide trapping, ExbD ceases to function. These results suggest that the snapshots of ExbB/D and MotA/B provided by recent cryo-EM structures do not yet capture the mechanism by which they utilize the PMF (50–52). We also exclude the overall importance of ExbD L132, first identified as an inactive L132Q missense mutation, showing that several different amino acids can functionally substitute at that position. Further, we identify an ExbD-specific cyclic peptide that, when added to *E. coli* exogenously, prevents binding of ExbD to its protein partners and inhibits the TonB system.

## RESULTS

### The negative charge and location of Asp25 in the ExbD transmembrane domain are indispensable for TonB system function

The ExbD transmembrane domain is predicted to consist of residues 23-43 (20). The similarity of the ExbD transmembrane domain to those of paralogues TolR and MotB suggested that ExbD D25 (D23 in TolR and D32 in MotB) might be important (48, 49). To investigate that possibility, Cys substitutions were engineered individually at all predicted ExbD transmembrane domain residues from 23 to 43 on plasmid pKP999 under a propionate-inducible promoter [(29); see Fig. 3A for the primary amino acid sequence]. Individual Cys substitution mutants were expressed at chromosomal levels in strain RA1045 (τ1*exbD,* τ1*tolQR*), where the paralogues TolQR were also removed and could no longer serve as a source of limited but detectable TonB system activity (28, 53–55). All Cys-substituted ExbD mutant proteins were proteolytically stable and, excluding the slower migration of ExbD D25C and ExbD P39C, migrated at the same position as wild type ExbD in immunoblots (Fig. 4, lower panel). The slower migration of D25C appeared to reflect the loss of the Asp residue since D25N and D25A substitutions also decrease ExbD mobility in sodium dodecyl sulfate (SDS) polyacrylamide gel electrophoresis (data not shown). A case where ExbD migrates more slowly because it appears to be chemically modified is considered below. Except for ExbD D25C, which was insensitive (tolerant) to all agents tested, the Cys substitutions of residues 23-43 supported sensitivity to colicin B, Ia, M and bacteriophage ϕ80 (Table 1). These TonB-system assays are the most sensitive, detecting the activity of as little as one TonB per bacterial cell (56).

**Figure 3.**
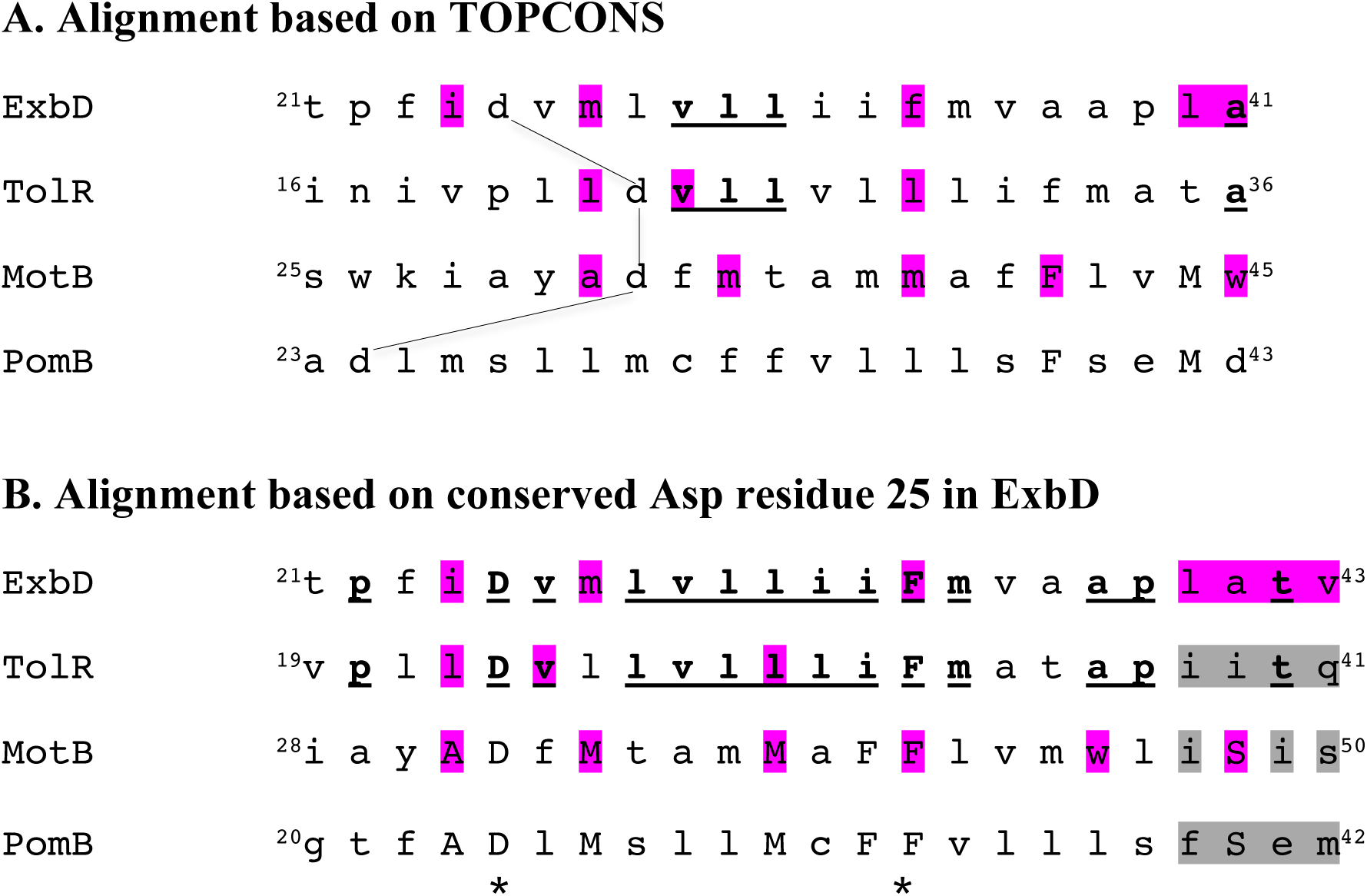
The most meaningful sequence comparisons among ExbD paralogues are based on the conserved Asp residue in the transmembrane domains. A slightly expanded number of possible transmembrane domain residues are considered here to encompass the widest range of options. Alignment based on the conserved Asp residue in B. has been previously proposed (48). Residues conserved between ExbD and TolR are indicated in bold underline in A and B. Residues conserved between MotB and PomB are in upper case in both A and B. Residues conserved in all four proteins in B. are indicated by an asterisk *. Residues which, as Cys substitutions, cross-link strongly with oxidative catalysis (I_2_ or cu-*o*- phenanthroline) are highlighted in magenta (46, 47). Cys substitutions in PomB have not been systematically analyzed for cross-links with I_2_ catalysis (121). Residues not evaluated as Cys substitutions, but that would be interesting to examine in the context of the results from this study, are in gray (46, 47, 121).

**Figure 4.**
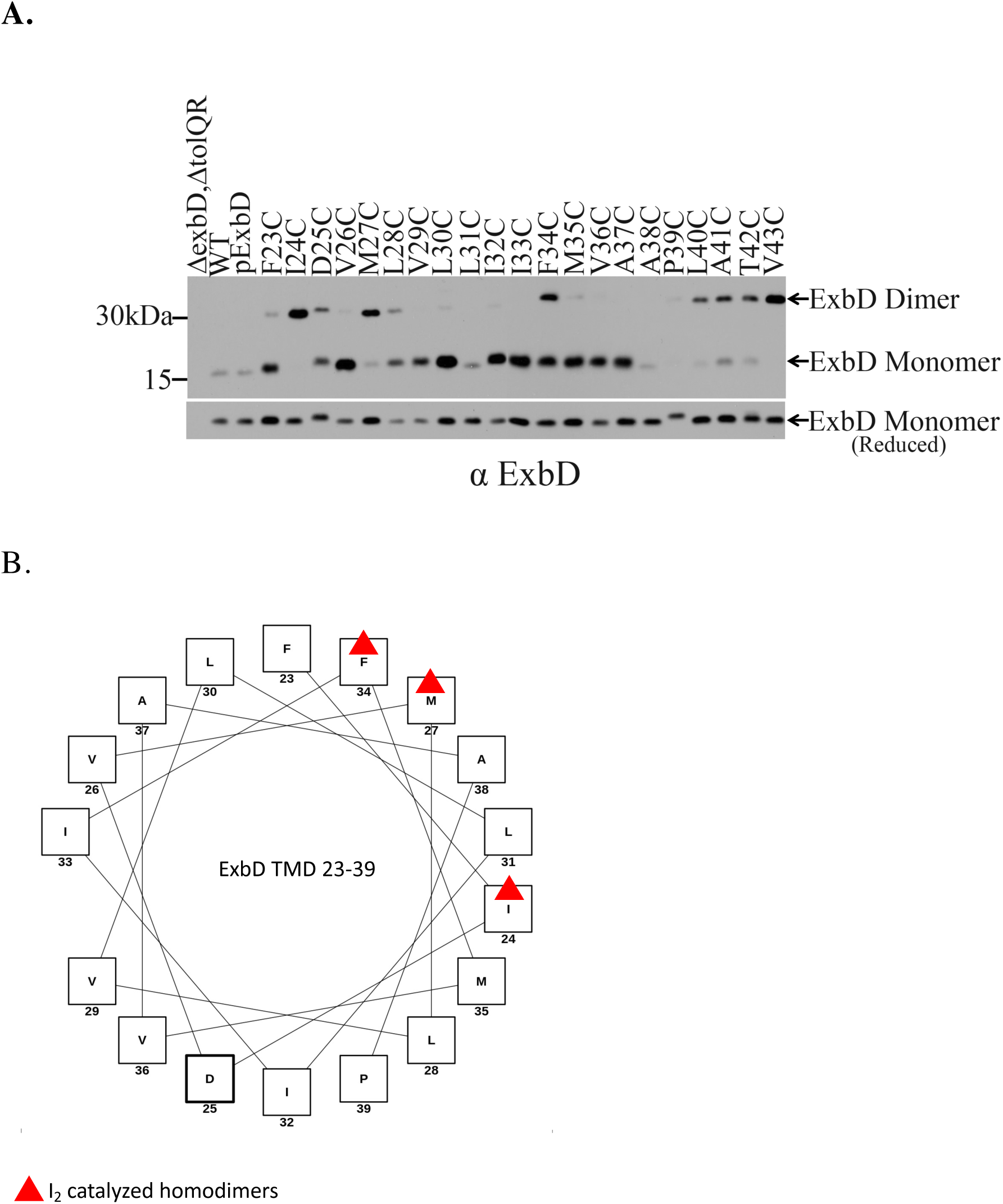
Oxidized Cys substitutions in the ExbD transmembrane domain define a helical homodimerization face. **A.** ExbD Cys substitutions in RA1045 (Δ*exbD*, Δ*tolQR*) were subjected to I_2_ catalysis as described in Materials and Methods. Divided samples were processed for immunoblotting with anti-ExbD antiserum on a non-reducing SDS 13% polyacrylamide gels. Strains and ExbD variants are indicated at the top of the immunoblot. The position of ExbD monomers and homodimers are indicated on the right with mass markers on the left. WT = W3110; pExbD = pKP999. The corresponding levels of ExbD variants resolved on a reducing SDS 13% polyacrylamide gel are indicated in the lower immunoblot. Equal numbers of cells were loaded in each lane. Two different immunoblots are shown with the partition occurring between ExbD I32C and I33C. **B.** Helical wheel diagram of residues 23-39 of the ExbD transmembrane domain. The most abundant complexes, formed by ExbD I24C, M27C and F34C, are labeled with red triangles and form a face on the helix. Residues 40-43 are not included in the helical wheel analysis since they represent a likely disordered region (see Fig. 5 results).

**Table 1.** Phenotypes of strains producing Cys-substituted ExbD proteins.

| **ExbD** | **ø80** | **colicin B** | **colicin Ia** | **colicin M** |
| --- | --- | --- | --- | --- |
| W3110 | 8, 9, 8 | 6, 6, 6 | 5, 5, 5 | 4, 4, 4 |
| pExbD | 8, 8, 8 | 5, 5, 5 | 5, 5, 5 | 4, 4, 4 |
| F23C | 6, 7, 6 | 4, 4, 5 | 5, 5, 4 | 4, 4, 4 |
| I24C | 6, 5, 6 | 4, 4, 4 | 5, 5, 5 | 3, 3, 3 |
| D25C | T, T, T | T, T, T | T, T, T | T, T, T |
| V26C | 7, 7, 7 | 5, 5, 5 | 5, 5, 5 | 4, 4, 4 |
| M27C | 7, 7, 7 | 5, 5, 5 | 5, 5, 5 | 4, 4, 4 |
| L28C | 7, 7, 7 | 4, 4, 4 | 5, 5, 6 | 4, 4, 4 |
| V29C | 8, 7, 7 | 5, 5, 5 | 5, 5, 5 | 3, 3, 3 |
| L30C | 6, 6, 6 | 4, 5, 4 | 5, 5, 5 | 3, 3, 4 |
| L31C | 6, 6, 6 | 5, 5, 5 | 5, 5, 5 | 3, 4, 4 |
| I32C | 6, 6, 6 | 5, 6, 5 | 5, 5, 5 | 4, 4, 4 |
| I33C | 7, 7, 7 | 4, 4, 4 | 5, 5, 5 | 3, 3, 3 |
| F34C | 8, 7, 7 | 5, 5, 5 | 4, 4, 4 | 3, 3, 3 |
| M35C | 7, 7, 7 | 4, 4, 5 | 5, 5, 4 | 3, 3, 3 |
| V36C | 8, 7, 7 | 5, 5, 5 | 4, 5, 4 | 3, 3, 3 |
| A37C | 8, 8, 8 | 5, 5, 5 | 5, 4, 5 | 4, 4, 4 |
| A38C | 7, 7, 7 | 5, 5, 5 | 4, 4, 4 | 3, 3, 4 |
| P39C | 7, 7, 7 | 4, 3, 4 | 4, 4, 4 | 3, 3, 3 |
| L40C | 8, 8, 8 | 5, 5, 5 | 5, 5, 5 | 4, 4, 4 |
| A41C | 8, 7, 7 | 5, 5, 5 | 4, 5, 4 | 3, 3, 3 |
| T42C | 7, 7, 7 | 5, 5, 5 | 5, 5, 5 | 3, 3, 3 |
| V43C | 8, 7, 7 | 5, 5, 5 | 5, 4, 5 | 3, 3, 3 |

Parent plasmid pExbD is pKP999. Phenotypes of ExbD mutants were evaluated in RA1045
( $\Delta exbD$ , $\Delta tolQR$ ) using spot titer assays (56). Cultures with the strains expressing ExbD mutated proteins at chromosomal levels were plated on T-plates and spotted with fivefold serial dilutions of colicins and 10-fold serial dilutions of bacteriophage $\Phi 80$ . Values were recorded as the reciprocal of the highest dilution at which clearing of the lawn was evident after 18 h of incubation at 37°C. ‘T’ indicates tolerance (insensitivity).

To further test the importance of Asp25 in ExbD, all 19 other amino acids were substituted at that position. The conservative replacement ExbD D25E retained 5-10% wildtype activity in iron transport assays, which are the most discriminative of the TonB system assays, while the remaining 18 substitutions at Asp25 were completely insensitive in colicin and phage sensitivity assays, indicating that they supported no TonB system activity (56).

Relocation of His20 in the TonB transmembrane domain from position 20 to a higher turn of the spiral at position 24 supports detectable but decreased colicin and phage sensitivity (26). To test the importance of its position within the ExbD transmembrane domain, Asp25 was relocated to all other positions in the ExbD transmembrane domain from position 23 to 43 on a plasmid template where Asp25 was mutated to Ala. In contrast to the situation with TonB His20, all the ExbD Asp25 relocation mutants were completely insensitive to colicin and phage. Taken together these results indicated that the negative charge and location of ExbD Asp25 within the transmembrane domain were essential for its activity.

### Sequence comparisons with ExbD paralogues TolR and MotB

We had previously used the TOPCONS program to predict ExbB, TolQ, MotA and PomA transmembrane domain boundaries (57). In that study, TOPCONS provided a prediction that was congruent with a number of conserved residues within their transmembrane domains and led to a redefinition of boundaries for the last two TMDs of each protein (27). Here TOPCONS analysis of the individual protein sequences was less helpful, potentially due to divergence in the function of these proteins compared to ExbB and its paralogues.

The invariant and essential Asp residue found in ExbD, TolR, MotB and PomB transmembrane domains did not align well using TOPCONS: the majority of homologies between the paralogous proteins ExbD and TolR were missed; congruence between ExbD and MotB Cys substitutions that formed high levels of disulfide-linked complexes were not optimized; and homologies between MotB and PomB, both of which play similar roles in flagellar rotation, were not optimized (Fig 3A). Because ExbD appears to interact with the last two transmembrane domains of ExbB, it seemed likely that TolR/MotB/PomB proteins should interact similarly with TolQ/MotA/PomA (27). This appeared to be the case since the inconsistencies generated by the TOPCONS analysis were resolved when the transmembrane domains were aligned using the invariant Asp residues (Fig. 3B). Notably, the ExbD Cys substitutions that homodimerized most efficiently (described below) were now aligned with the MotB substitutions that homodimerized most efficiently.

### ExbD homodimerizes through residues 23-63 in vivo

We had previously identified residues in the region of residues 92-121 through which ExbD periplasmic domains homodimerized *in vivo* (36). To study the ability of the ExbD transmembrane domain to support homodimerization in the absence of its periplasmic domain, the DNA encoding ExbD residues V23-P63 was cloned into the TOXCAT vector, pccKan (58). The cloned region included the predicted ExbD transmembrane domain [(residues 23-43); (20)] and a subsequent unstructured domain involved in signal transduction [(residues 44-63) (59, 60)].

As part of the TOXCAT fusion protein, if residues 23-63 homodimerized they would mediate concomitant homodimerization of the amino terminal, cytoplasmically localized ToxR domain, thus activating the chloramphenicol acetyl transferase gene (CAT) and supporting increased resistance to chloramphenicol (CAM) compared to controls where the transmembrane domain does not homodimerize (GlpA compared to GlpA G83I in the TOXCAT vector). Activity of carboxy terminally fused periplasmic maltose binding protein (MBP) would serve as a control to demonstrate that the inserted ExbD residues had translocated MBP to the periplasmic space.

To test homodimerization by ExbD residues 23-63, plasmid vector pccKan, plasmid pccGlpA, plasmid pccGlpA G83I and pKP523 [pccExbD (*toxR*-*exbD*_23-63_-*malE*)] were transformed into strain NT326 [*araD*139, Δ(*argF-lac*)_U169_, Δ*malE444, recA*1, *thi*, *strR* (61)] and KP1448 (NT326, *exbB*::Tn*10*). Resistance to various concentrations of CAM, from 34 *µ*g/ml to 1.0 mg/ml was measured by growth on plates. In NT326, both GlpA-TOXCAT and ExbD_23-63_-TOXCAT supported strong growth on 125 *µ*g/ml CAM and poor growth at 500 *µ*g/ml. The GlpAG83I-TOXCAT mutant control supported poor growth at 125 *µ*g/ml and no growth at 500 *µ*g/ml (Table 2). In KP1448, where competition by wild-type ExbD transmembrane domains was eliminated, ExbD_23-63_-TOXCAT and GlpA-TOXCAT again supported equally robust growth on 0.125 *µ*g/ml CAM. Again, there was little growth of the negative control, GlpA G83I-TOXCAT, at that concentration (Table 2). The three TOX-CAT constructs containing GlpA, GlpA G83I, or ExbD_23-63_ residues equally supported growth with maltose as sole carbon source and had equivalent levels of protein present on anti-MBP immunoblots (data not shown), indicating that they were equally expressed and exported to the cytoplasmic membrane. We concluded that ExbD residues 23-63, which included the predicted transmembrane domain, had an intrinsic ability to form homodimers *in vivo* in both the presence and absence of ExbB.

**Table 2.** ExbD residues 23-63 form homodimers independently of ExbB.

| Plasmid | <sup>a</sup>34 | 62.5 | 125 | 250 | 500 | 1000 |
| --- | --- | --- | --- | --- | --- | --- |
| <sup>b</sup>WT/pccKan vector | - | - | - | - | - | - |
| WT/pccGlpA | <sup>c</sup>n.d. | 4+ | 4+ | 4+ | 1+ | - |
| WT/pccGlpA G83I | n.d. | 2+ | +/- | - | - | - |
| WT/pccExbD<sub>23-63</sub> | n.d. | 4+ | 3+ | 2+ | +/- | - |
| <sup>d</sup>*exbBD*/pccKan vector | - | - | - | - | - | - |
| *exbBD*/pccGlpA | 4+ | 4+ | 4+ | 3+ | 1+ | - |
| *exbBD*/pccGlpA G83I | 3+ | 2+ | 1+ |  |  |  |
| *exbBD*/pccExbD<sub>23-63</sub> | 4+ | 4+ | 4+ | 3+ | 1+ |  |

<sup>a</sup> plate concentration of chloramphenicol (μg/ml)
<sup>b</sup> strain NT326 is wild-type for *exbB*: [*araD*139, Δ(*argF-lac*)<sub>U169</sub>, Δ*malE*444, *recA*1, *thi*, *strR* (61)]
<sup>c</sup> growth; n.d. = not determined
<sup>d</sup> strain NT326, made *exbB*::Tn10 by P1vir transduction (KP1448). The Tn10 insertion in *exbB* is polar on downstream *exbD* expression (119).

### Identification of a face through which ExbD transmembrane domains homodimerize

To determine if homodimers formed through the predicted ExbD transmembrane domain, disulfide-mediated homodimerization of ExbD Cys-substitutions at residues 23-43 expressed at chromosomal levels was evaluated using iodine (I_2_) catalysis, a technique previously used to study homodimerization of TolR and MotB transmembrane domains (46, 47). In Fig. 4A, several of the Cys substitutions resulted in bands on non-reducing anti-ExbD immunoblots that, at ∼ 31 kDa, were twice the mass of monomeric ExbD (∼15 kDa) (45). The ∼ 31 kDa complex was absent from an ExbD deletion strain (RA1045, Δ*exbD,* Δ*tolQR*) and from wild-type (Cys-less) ExbD samples (WT and pExbD), indicating that it represented a disulfide-linked ExbD homodimer. The efficiency of dimer formation varied with the position of the Cys residue. The most abundant dimers were formed by residues I24C, M27C, F34C, with a periodicity of ∼3-4 amino acids (and a gap at 30-31C), consistent with an α-helical arrangement of each ExbD transmembrane domain and suggesting that they homodimerize through one face (Fig. 4B).

Similar results are seen for paralogues MotB and TolQ (46, 47). TolR, MotB, and as we demonstrated above, ExbD, all carry a functionally important Asp residue in their single transmembrane domain at positions 23, 32, and 25 respectively (47, 49). Further similarity was demonstrated by ExbD I24C, which existed entirely in homodimeric form, with ExbD D25C existing mostly in monomeric form in this assay (Fig. 4A). A similar relationship was seen with corresponding MotB transmembrane domain Cys substitutions at A31C and D32C (46), suggesting that the transmembrane domains homodimerize similarly. In contrast to a predicted α-helical configuration, each of the ExbD Cys substitutions from L40C-V43C formed homodimers with little to no monomer. It was not clear if the I_2_ oxidant or ExbB was needed to form any of the complexes.

Although all the ExbD mutants were expressed at near chromosomal levels as determined by reducing SDS polyacrylamide gel electrophoresis (Fig. 4A, lower), there was differential antibody detection of both monomers and homodimers from the same samples immunoblotted under non-reducing conditions (Fig. 4A, upper). For example, V26C monomer was detected at just slightly less than wild-type monomer ExbD levels on the reducing immunoblot (Fig. 4A, lower) but was much more strongly detected compared to the wild-type ExbD on the non-reducing immunoblot (Fig. 4A, upper) and did not form homodimers. In contrast, ExbD P39C appeared to be undetectable on the non-reducing immunoblot and was present at wild-type chromosomal levels on the reducing immunoblot. The differential antibody detection suggested that there was structural variation among ExbD Cys substitutions under non-reducing conditions, even in the presence of SDS.

### ExbD transmembrane domain Cys substitutions 23-39 require oxidant for homodimerization whereas Cys residues at 40-43 do not

To determine if oxidant was required for transmembrane domain homodimerization, the experiment in Fig. 4A was repeated but without the use of I_2_ (Fig. 5). Near the cytoplasmic side of the predicted transmembrane domain, I24C spontaneously formed only modest levels of homodimers in contrast to the high level I24C formed when I_2_ was present, thus placing it within the transmembrane domain. In contrast, ExbD F23C appeared to be on the transmembrane domain boundary because it formed similarly low levels of homodimers with and without I_2_ (compare Figs. 4A and 5). Consistent with an apparent transmembrane domain location, none of the residues from D25C through P39C spontaneously formed significant levels of homodimers in the absence of I_2_.

**Figure 5.**
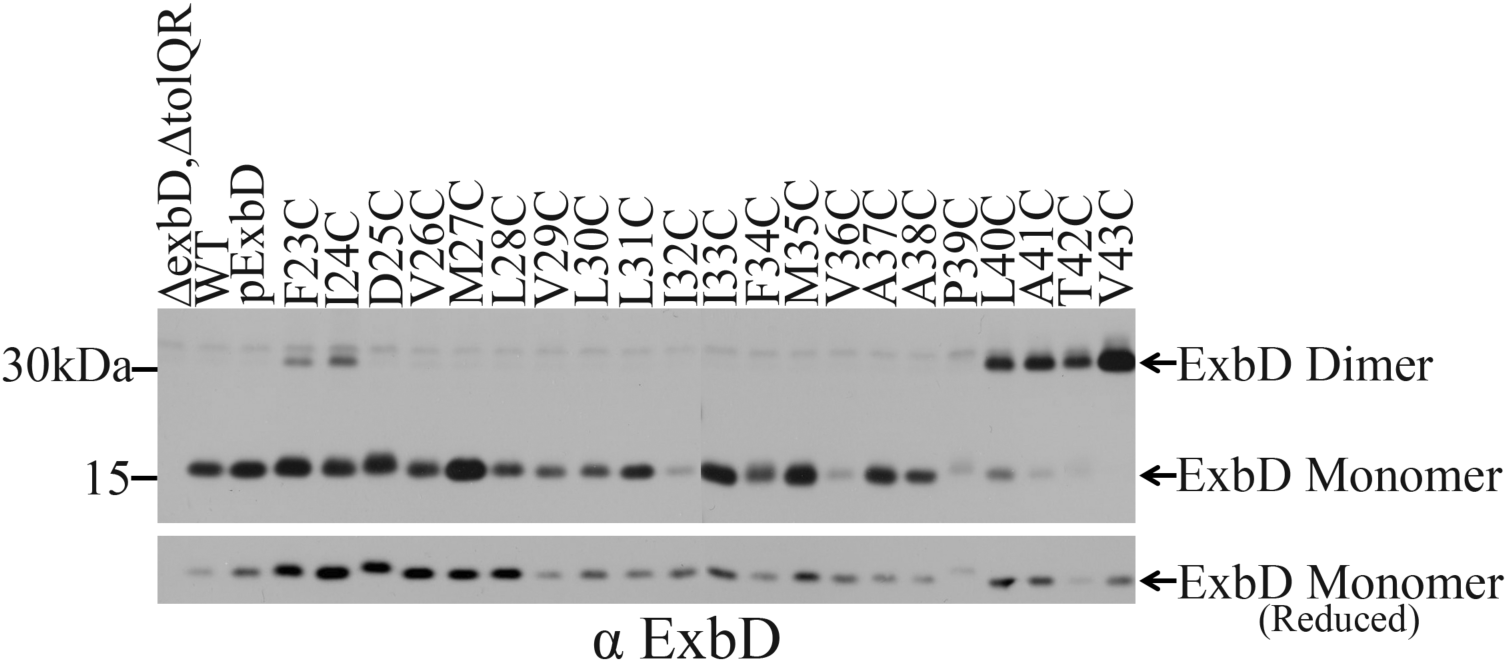
Spontaneous homodimer formation through ExbD L40C, A41C, T42C, and V43C suggests they are separate from the helical transmembrane domain. Divided samples of ExbD Cys substitutions in RA1045 (Δ*exbD*, Δ*tolQR*) prepared in the absence of I_2_ were processed for immunoblotting with anti-ExbD antiserum on non-reducing SDS 13% polyacrylamide gels as described in Materials and Methods (upper). Strains and ExbD variants are indicated at the top of the immunoblot. The position of ExbD monomers and homodimers are indicated on the right with mass markers on the left. WT = W3110; pExbD = pKP999. Two different immunoblots are shown with the partition occurring between ExbD I32C and I33C. The corresponding levels of ExbD variants resolved on a reducing SDS 13% polyacrylamide gel are indicated in the lower immunoblot. Equal numbers of cells were loaded in each lane.

In contrast, residues from ExbD L40C-V43C each spontaneously formed high levels of homodimers in the absence of I_2_, suggesting they were not part of the transmembrane α-helix and likely constituted a region of intrinsic disorder (compare Figs. 4A and 5).

### Trapping ExbD homodimers through T42C or V43C largely inactivates the TonB system

Because ExbD T42C and V43C resulted in almost 100% homodimer formation in the absence of I_2_ oxidant in Fig. 5, it was possible to determine whether dynamic movement of those residues was required for TonB system activity. We assayed the ability of plasmids expressing each variant to support TonB-dependent iron transport with or without the reducing agent dithiothreitol (DTT) and immunoblotted those same cultures to determine relative steady state levels of monomer and homodimer. As seen previously, both ExbD T42C and V43C were nearly 100% homodimerized in the absence of DTT (Fig. 6A, upper panel). All ExbD Cys substitutions were expressed at chromosomally encoded levels (Fig. 6A, lower panel).

**Figure 6:**
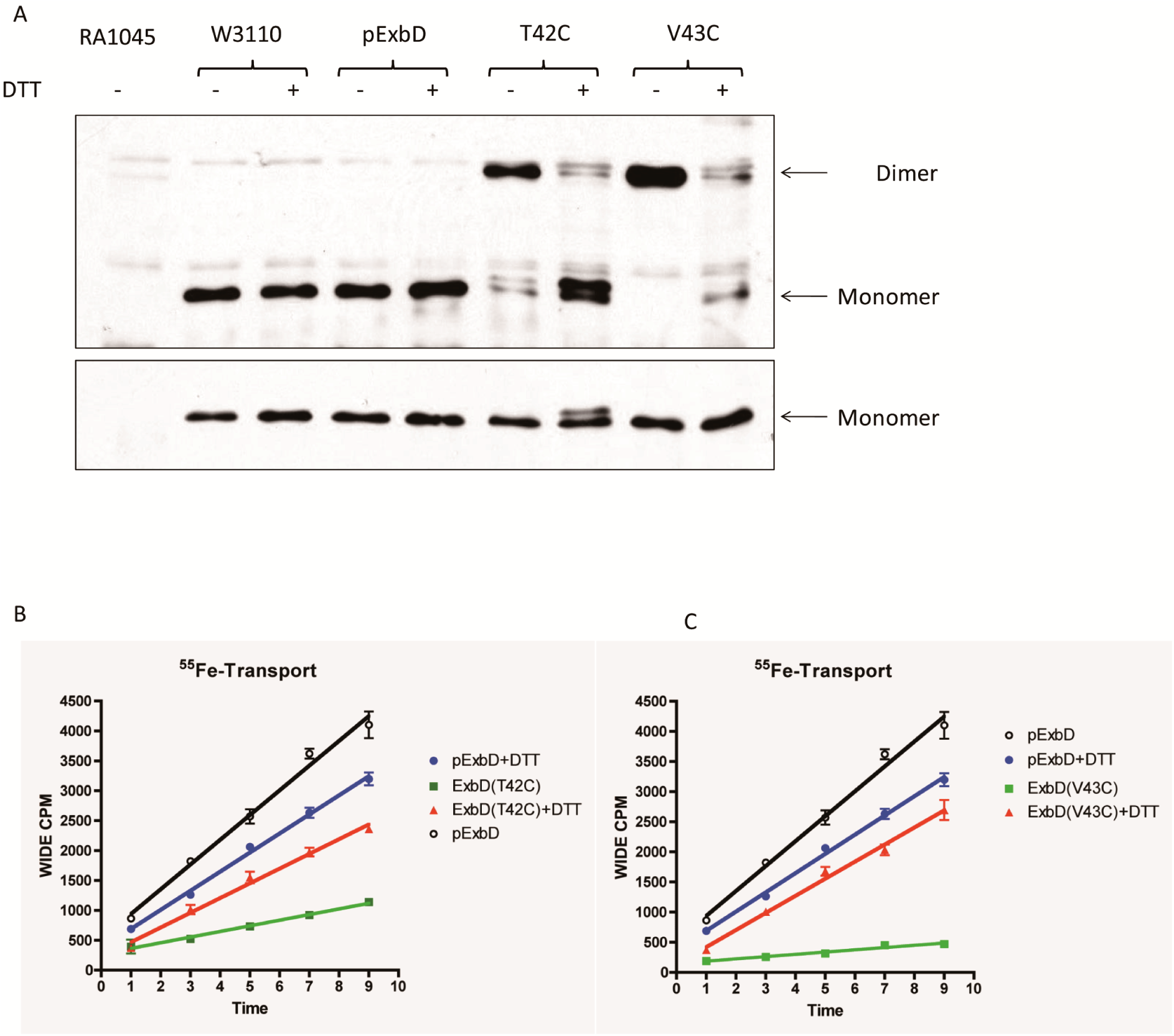
DTT treatment releases ExbD disulfide-trapped through residues T42C and V43C to support TonB-dependent iron transport. **A.** Immunoblot of a non-reducing SDS 13% polyacrylamide gel with anti-ExbD antiserum of the samples evaluated for iron transport. Identity of strains and ExbD variants is indicated at the top. Presence (+) or absence (-) of 2 mM DTT during the logarithmic phase of growth is indicated. The lower panel is an immunoblot of a reducing SDS 13% polyacrylamide gel to determine relative steady state levels of the same samples. Positions of ExbD monomer or dimer are indicated on the right. Equal numbers of cells were loaded in each lane. **B.** Initial rates of iron transport for RA1045 (Δ*exbD*, Δ*tolQR*) cultures with ExbD plasmids, both with and without DTT. Left panel evaluates plasmids expressing ExbD T42C; right panel evaluates plasmids expressing ExbD V43C. See Table 3 for calculation and comparison of initial rates.

**Table 3.** ExbD T42C and V43C disulfide-linked homodimers are impaired in TonB system-mediated iron transport.

|  | % rate relative to pExbD<br>without DTT | % rate relative to<br>pExbD with DTT |
| --- | --- | --- |
| **RA1045** | 0.15% |  |
| **W3110** | 121% |  |
| **pExbD** | 100% | 100% |
| **pExbD L40C** | 101.9% |  |
| **pExbD A41C** | 103.6% |  |
| **pExbD T42C** | 22.9% | 77% |
| **pExbD V43C** | 9.2% | 89% |

RA1045 is [ $\Delta exbD$ , $\Delta tolQR$ ]; (54)]; W3110 is wild-type *E. coli* K12 (111); pExbD is pKP999 (29).

In Fig. 6B, DTT had an inhibitory effect, decreasing the initial transport rate of the wild-type control to 77% of the rate seen in the absence of DTT. We compensated for the fact that DTT was somewhat inhibitory to wild-type transport by calculating the mutant transport rates as percentiles of the wild-type transport rates. The effect of DTT on the previously trapped inactive homodimers of ExbDT42C and ExbDV43C, was to release them as monomers, thus restoring activity (Fig 6A, upper panel). Transport supported by trapped ExbD V43C was restored from 9% of wild-type-DTT to 89% of wild-type +DTT (Fig. 6C; Table 3). Similarly, DTT restored transport supported by ExbD T42C from 23% of wild-type-DTT to 77% of wild-type +DTT. These results indicated that inactivity of the disulfide-linked homodimers was due to restriction of required conformational changes rather than the inability of Cys to functionally replace Val or Thr. The results were not contradictory with the spot titer data from Table 1 where strains expressing ExbD T42C or V43C were sensitive to colicins and phage due to the presence of residual monomer. The highly sensitive colicin spot titers detect activity from as little as 1 molecule of active TonB (56, 62).

The results were consistent with the existence of an energy transduction cycle where, if ExbD is trapped in a homodimeric configuration such as Stage I or II, it cannot progress through the cycle (Fig. 2). Residues L40-V43 are contiguous with a region of intrinsic disorder from 44-63 that dynamically interacts with a second ExbD, with ExbB and with TonB *in vivo* (35, 59, 60).

### ExbD T42C appears to undergo increased modification in the absence of ExbB

Data presented in Fig. 6A (lower panel) suggested that monomeric ExbD T42C might be chemically modified, because while a majority of ExbD T42C was found migrating at the same position as wild-type ExbD on immunoblots of reducing SDS polyacrylamide gels, a smaller amount appeared to migrate at a slightly higher apparent mass. To determine the parameters that affected the potential modification, the migration of ExbD T42C was evaluated on immunoblots of strains lacking ExbD, both ExbB and ExbD, or both ExbD and TonB (Fig. 7). As seen in Fig. 6A, both unmodified and modified ExbD T42C were again observed, with ∼ 2/3 in the unmodified form (Fig. 7, lane 6). The absence of ExbB (and paralogue TolQ) converted all the ExbD T42C into the higher mass form (Fig. 7, lane 7). In the absence of TonB, the relative proportion of the two forms was reversed relative to the wild-type situation, with ∼2/3 of the ExbD T42C found at the position of the higher mass form (Fig. 7, lane 8). It is not clear whether the modification has any *in vivo* relevance for the mechanism of TonB-dependent energy transduction. At the very least it provides evidence of *in vivo* interactions between ExbD and ExbB. It is included here for future researchers. Apparent chemical modification of full-length wild-type ExbD also occurs when an inhibitory ExbD peptide (residues 44-63) is expressed endogenously, resulting in the appearance of a higher mass ExbD species (63).

**Figure 7.**
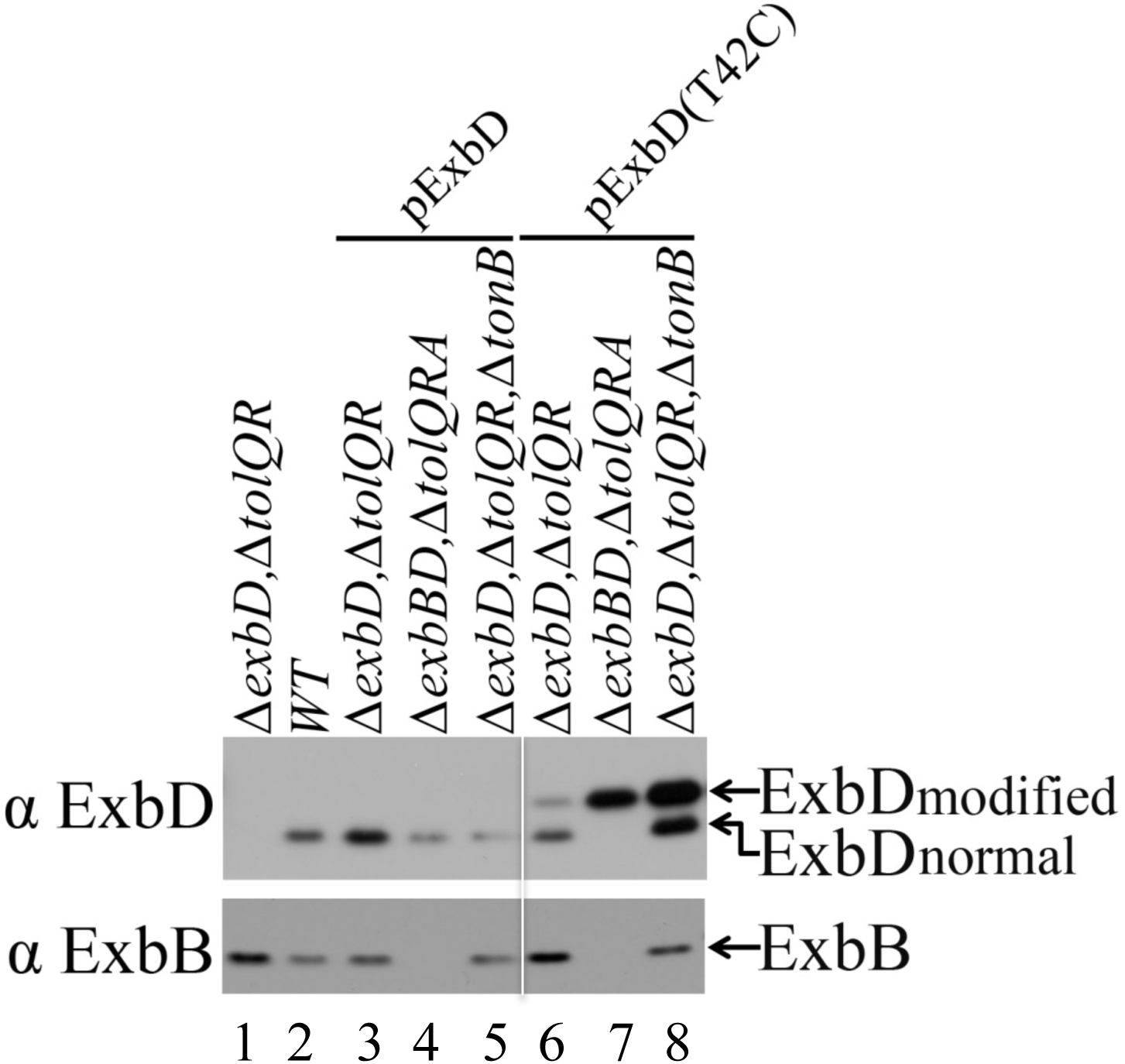
Apparent *in vivo* modification of ExbDT42C occurs most prominently in the absence of ExbB. Wild-type plasmid-encoded ExbD (pKP999) or ExbDT42C (pKP1392) were grown as described in Materials and Methods and immunoblotted under reducing conditions with anti-ExbD antibodies (upper panel) or anti-ExbB antibodies (lower panel). The positions of normal and modified ExbD and of ExbB are indicated on the right. Strain genotypes are listed at the top of the immunoblot; WT = W3110. RA1045 is (Δ*exbD*, Δ*tolQR*); RA1017 is (Δ*exbBD*, Δ*tolQRA*); KP1509 is (Δ*exbD*, Δ*tolQR, ΔtonB*). Lane 1, RA1045; lane 2, W3110; lane 3, RA1045/pKP999; lane 4, RA1017/pKP999; lane 5, KP1509/pKP999; lane 6, RA1045/pKP1392; lane 7, RA1017/pKP1392; lane 8, KP1509/pKP1392. Equal numbers of cells were loaded in each lane. The immunoblots shown for each antibody are from the same exposure with non-relevant lanes removed (indicated by the white line). Amounts of propionate inducer added were: lane 1, none; lane 2, none, lane 3, 0.25 mM; lane 4, 3.0 mM, lane 5, none; lane 6, 0.018 mM; lane 7, 0.02 mM; lane 8, none.

### Transmembrane domain Cys residues require ExbB for homodimerization whereas those of the potentially disordered domain do not

Because the ExbD_23-63_-TOXCAT fusion protein homodimerized in the absence of ExbB, we tested the role of ExbB in ExbD homodimerization through its transmembrane domain by expressing each of the Cys-substituted proteins in RA1017 (*11exbBD, 11tolQRA*). Unexpectedly, in light of the TOXCAT results, the absence of ExbB prevented the previously detected I_2_-dependent disulfide-linked dimerization through ExbD substitutions F23C-P39C, even though I_2_ was present (compare Figs. 4 and 8). This finding notably contrasts with MotB transmembrane domain homodimer formation through engineered Cys substitutions, which does not depend on MotA (46).

**Figure 8.**
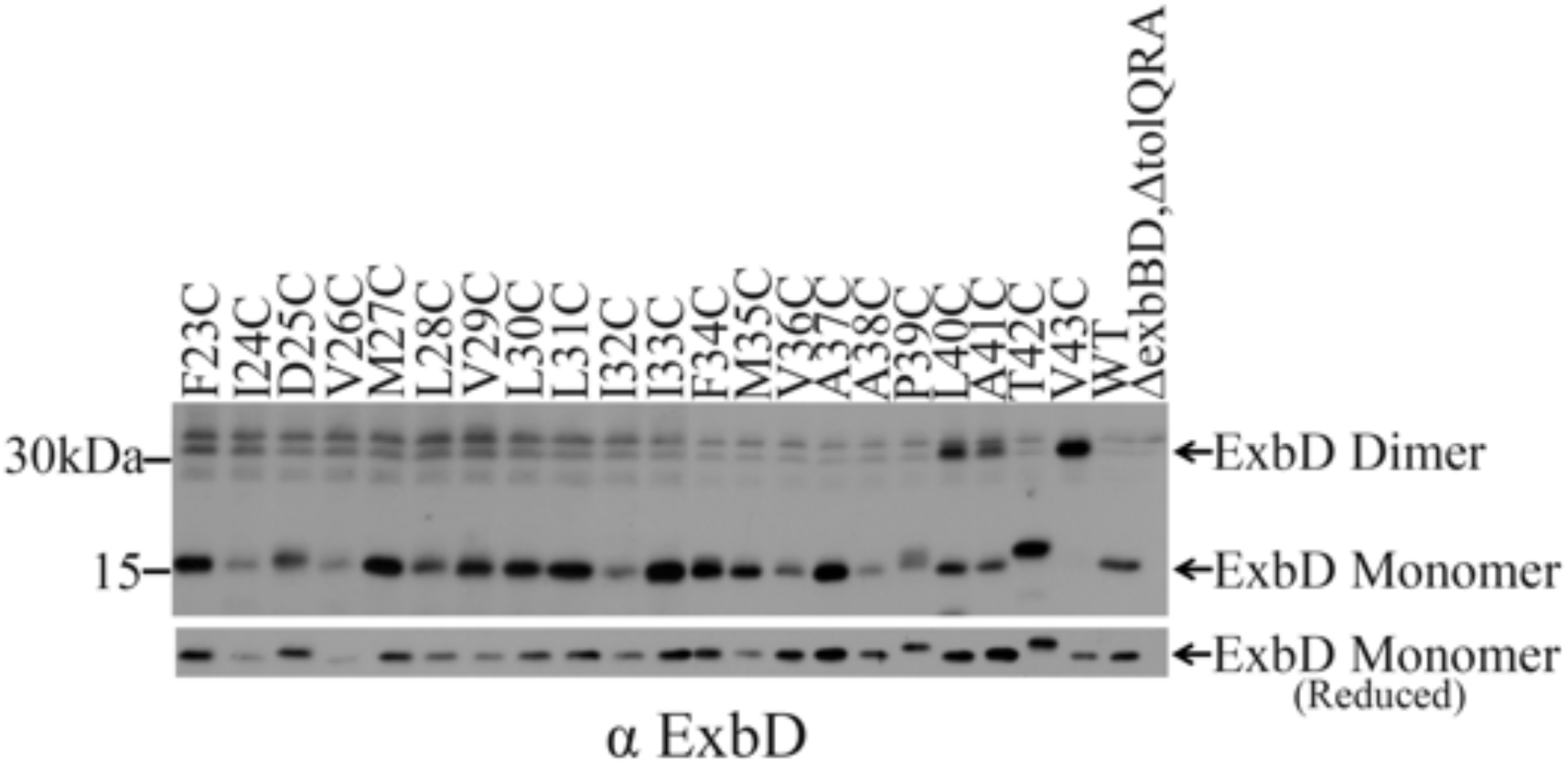
ExbB is required for homodimerization of full-length ExbD through its transmembrane domain. ExbD Cys substitutions in RA1017 (11*exbBD,* 11*tolQRA*) were subjected to I_2_ catalysis as described in Materials and Methods. Divided samples were processed for immunoblotting with anti-ExbD antiserum on non-reducing SDS 13% polyacrylamide gels as described in Materials and Methods (upper). Strains and ExbD variants are indicated at the top of the immunoblot. The position of ExbD monomers and homodimers are indicated on the right with mass markers on the left. WT = W3110. Two different immunoblots are shown with the partition occurring between ExbD I33C and F34C. ExbB T42C did not homodimerize, likely because it was chemically modified in the absence of ExbB (see Fig. 7). The corresponding steady state levels of ExbD variants resolved on a reducing SDS 13% polyacrylamide gel are indicated in the lower immunoblot. Equal numbers of cells were loaded in each lane.

In contrast, even without ExbB, residues contiguous to the transmembrane domain, L40C, A41C and V43C, formed homodimers, with V43C again achieving nearly 100% dimer formation (Fig. 8). The results suggested that because two ExbD molecules were in sufficient proximity to homodimerize through an adjacent region of likely intrinsic disorder, it seemed likely that the transmembrane domains also interacted, but in a way not captured by disulfide cross-linking in the presence of I_2_. In addition, these results suggested that the source of ExbB-independent homodimerization by the ExbD_23-63_-TOXCAT fusion protein resided in the disordered domain.

In the absence of ExbB, ExbD T42C formed essentially no homodimers, suggesting either that the Cys substitution was the site of chemical modification observed in Fig. 7 or that without ExbB, two T42C residues were no longer in proximity for disulfide formation. The latter explanation was excluded because A41C and V43C on either side of T42C formed disulfide-linked homodimers (Fig. 8). We therefore conclude that chemical modification occurred at ExbD T42C in the absence of ExbB.

### *In vivo* photo-cross-linking suggests that ExbD transmembrane domains rotate relative to each other and relative to ExbB

Disulfide linkage of ExbD Cys substitutions trapped transmembrane domains in a static homodimeric state where the most abundant cross-linking occurred between residues on an apparently helical face (Fig. 4A). Nonetheless, there was minor cross-linking that occurred through two ExbD D25C residues on an opposing face of that helix, consistent with the idea that the transmembrane domain helices might rotate relative to one another. Because the homodimerized transmembrane domains of TolR, which can functionally substitute for ExbD in the TonB system, rotate relative to one another *in vivo* (47), we wanted to determine if that was also true for ExbD. Rather than disulfide cross-linking, we used *in vivo* photo-cross-linking by the reagent *p*-benzoyl-l-phenylalanine (pBpa), which inserted efficiently at each amber codon for T21, V23, I24, D25, M27, V29, I32, D34, M35, and A37 (Fig. 9A, lower panel; monomeric ExbD).

**Figure 9.**
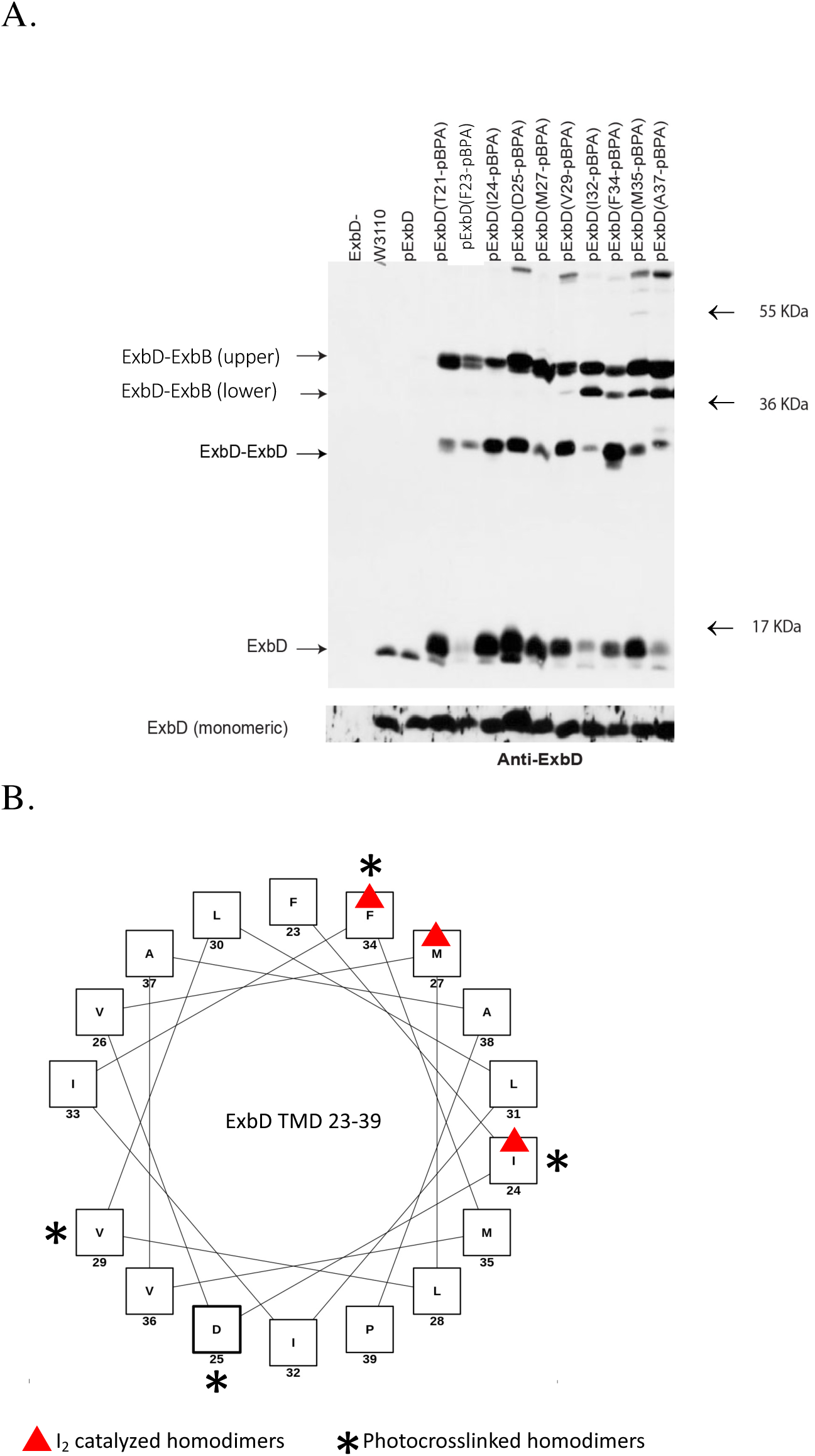
*In vivo* photo-cross-linking suggests that the ExbD helices rotate with respect to one another. **A.** ExbD with pBpa incorporated at various codons in the transmembrane domain was subjected to *in vivo* photo-cross-linking as described in Materials and Methods, resolved on an SDS 13% polyacrylamide gel and immunoblotted with anti-ExbD antisera. Strains and substituted ExbD residues are indicated at the top of the upper immunoblot. Strain labeled ExbD-is RA1045 (Δ*exbD*, Δ*tolQR*); pExbD is pKP999; W3110 is wild-type *E. coli.* The position of ExbD monomers, homodimers, and two different previously identified ExbD-ExbB homodimers (59), are indicated on the left with mass markers on the right. Prior to treatment of the samples in A. with ultraviolet light, the level of steady state monomeric ExbD expression and incorporation of pBpA were determined by processing the same volume of sample from A. for immunoblotting with anti-ExbD antiserum on reducing SDS 13% polyacrylamide gels shown in the lower immunoblot. Equal numbers of cells were loaded in each lane. **B.** Helical wheel diagram of residues 23-39 of the ExbD transmembrane domain. The most abundant homodimers formed by *in vivo* photo-cross-linking occurred through ExbD I24pBpa, D25pBpa, V29pBpa, and F34pBpa and are labeled with asterisks. The most abundant disulfide-linked homodimers, formed by ExbD I24C, M27C and F34C, are labeled with red triangles as in Fig. 4. Residues 40-43 are not included in the helical wheel analysis since they represent a likely disordered region (see Fig. 5 results).

In the presence of ultraviolet light, the ExbD pBpa monomers were pulled into three higher mass complexes on immunoblots: ExbD homodimers and two ExbD-ExbB complexes with different apparent masses (termed “upper” and “lower”) characterized previously [Fig. 9A, upper panel (59)]. Each of the ExbD pBpa substitutions formed homodimers and to different extents. Abundant homodimers were formed by I24pBpa and F34pBpa, two sites that, as Cys substitutions, were part of the proposed α-helical dimerization face (compare Figs. 4 and 9A). In addition, equally abundant homodimers were formed by D25pBpa and V29pBpa, two residues on the opposite side of that face (Fig. 9 A and B). Taken together, the results indicated that each of the transmembrane domain pBpa substitutions had access to a second, homodimerized ExbD, most likely through its transmembrane domain. To achieve contacts through residues on opposite faces of the α-helix, the transmembrane domain must have moved relative to a second ExbD, possibly by rotation. Because all the transmembrane domain ExbD pBpa substitutions formed complexes with ExbB (upper), they may have rotated relative to ExbB as well. Half of the pBpa-substituted ExbDs also formed complexes with ExbB (lower) (Fig. 9A).

### ExbD L132 plays only a structural role in the TonB system

The Braun lab isolated the first inactive missense mutation in the ExbD periplasmic domain, L132Q (43). It stalls TonB at Stage I of the energy transduction cycle and, unlike wild-type ExbD, does not formaldehyde cross-link to TonB *in vivo* [Fig. 2; (42)]. Deletion of the last 10 residues of ExbD (Δ132-141) render it inactive, possibly due to the loss of L132 (35). To determine if L132 was an essential residue like D25 in the ExbD transmembrane domain, we engineered substitutions of all 19 amino acid residues at position L132 in pKP999. The relative activity of the ExbD variants was assessed by ability to complement the RA1045 Δ*exbD* mutation in a growth assay in M9 minimal medium, where iron is limiting and only the TonB+ strains grow. As shown in Table 4, L132 is not an essential residue since it could be substituted with Cys, Val, Ile, Met, Phe, and Trp and still retain full activity. It seems likely that the Gln substitution in L132Q inactivated the TonB system by preventing ExbD interaction with TonB through steric hindrance.

**Table 4.**
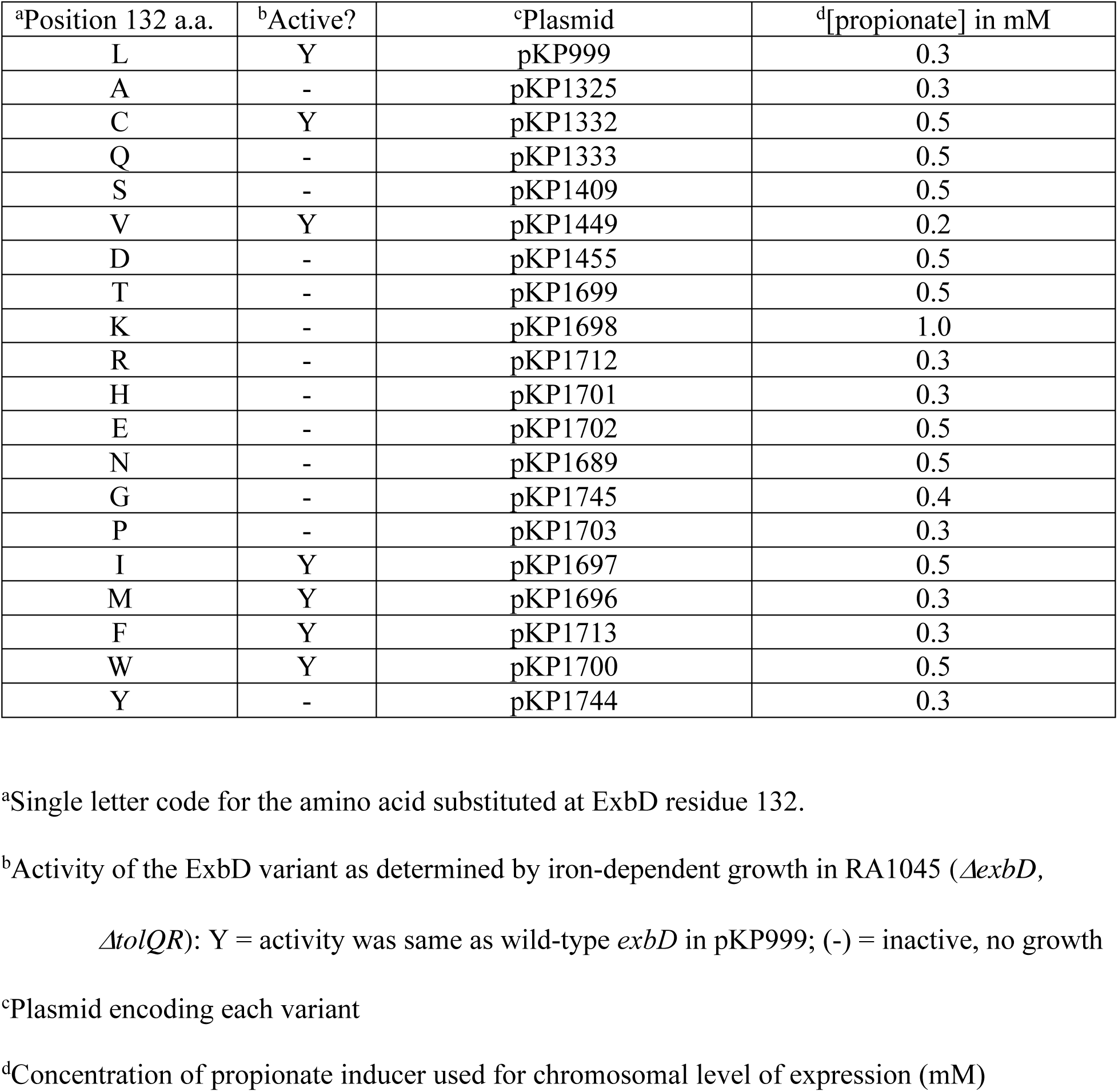
ExbD L132 is not an essential residue

### A cyclic peptide derived from ExbD inhibits TonB system activity and prevents formaldehyde-specific TonB-ExbB and TonB-ExbD cross-linked complexes in vivo

The TonB system is a virulence factor for many Gram-negative bacteria (9). The periplasmic domains of two key proteins in the system, ExbD and TonB, and their required interaction for subsequent active transport through outer membrane TBDTs make it an appealing target for novel antibiotic development. In general, antibiotics that inhibit periplasmic activities need possess only the hydrophilic chemical characteristics that allow diffusion through outer membrane porins to access their targets, a distinct advantage over those that must additionally cross the hydrophobic cytoplasmic membrane (64).

Periplasmic ExbD residues 44-63 are disordered and thus a likely site for interaction with other proteins (60, 65). We began to identify inhibitory regions of ExbD using endogenous secretion of promising small peptides into the periplasmic space. Secretion of a disordered region peptide (residues 44-63) inhibits TonB-dependent iron transport by inhibiting an essential ExbD-TonB interaction, thus establishing a proof of principle (63).

In this work we targeted interactions between ExbD and TonB periplasmic domains using exogenously added peptides. Cyclic rather than linear peptides were chosen to decrease the likelihood of proteolytic degradation by periplasmic proteases. None of the peptides contained cleavage sites for the outer membrane protein, OmpT (66). Inhibitory cyclic peptides were designed based on three regions of ExbD that are essential for TonB system activity *in vivo* [Fig. 10; (35, 36, 44)]: Region 1, residues 44-63 from the ExbD periplasmic disordered region adjacent to the transmembrane domain, an essential motif required for the ExbD-TonB PMF-dependent interaction (59); Region 2, residues 92-121 that are required for *in vivo* formaldehyde cross-linking of all known ExbD complexes (ExbD-ExbD, ExbD-TonB, and ExbD-ExbB) (35); and Region 3, the region surrounding ExbD residue, L132, where a single substitution, L132Q, completely abolishes activity (43).

**Figure 10:**
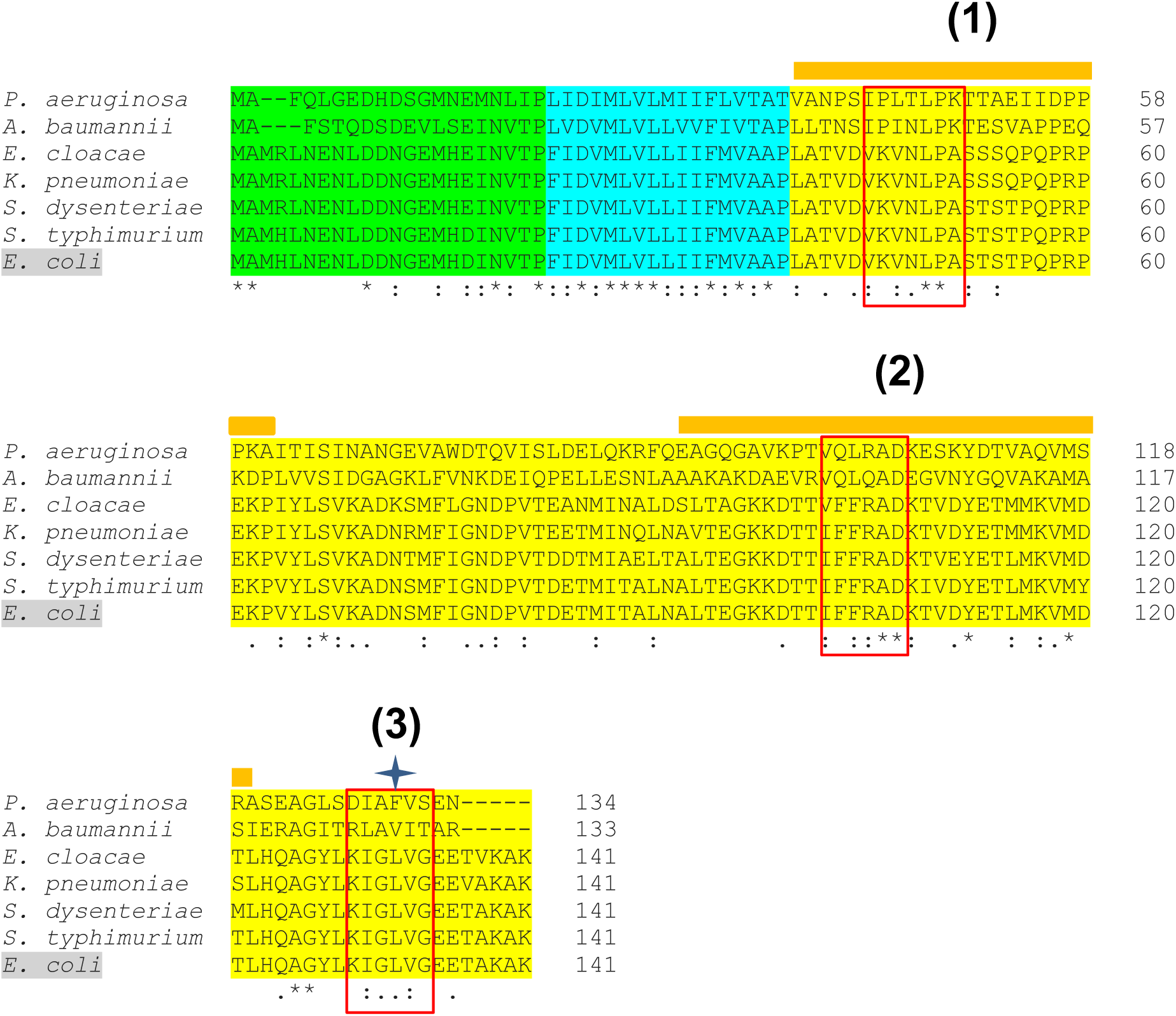
ExbD sequences chosen as cyclic peptide templates are semi-conserved among several ESKAPE pathogens. Based on the *E. coli* K12 primary amino acid sequence, the ExbD cytoplasmic domain is highlighted in green, the transmembrane domain is highlighted in blue, and the periplasmic domain is highlighted in yellow. The *E. coli K12 exbD* DNA sequence is identified by the gray highlight in the sequence alignment. The selected ExbD regions from which synthetic cyclic peptides were designed based on *E. coli* K12 sequences are boxed in red. The asterisks (*) below the alignment identify complete residue conservation; hyphens (:) identify conserved residue properties; periods (.) identify conserved similar residue properties. The orange bar labeled (**1**) above the alignment identifies the ExbD disordered domain in *E. coli* K12—residues 40-43 from this study plus residues 44-63 from (60). The orange bar labeled (**2**) above the alignment identifies the ExbD residues 92-121 in *E. coli* K12 that are necessary for all known ExbD formaldehyde cross-linked complexes (35). The blue star labeled (**3**) above the alignment identifies the L132 residue shown in this work to be relatively unimportant. The accession numbers are *P. aeruginosa* (AAF04084.1); *A. baumannii* (AOX72224.1); *E. cloacae* (ESS57504.1); *K. pneumoniae* (CDO12443.1); *S. dysenteriae* (ABB63085.1); *S. typhimurium* (TKE83773.1); *E. coli K12* (AAA69172.1). Alignment was generated using Clustal-Omega (1.2.4) multiple sequence alignment (http://www.clustal.org/omega).

The cyclic peptides from Region 1 of ExbD were the motifs V-K-V-N-L-P and V-K-V-N-L-P-A, where the latter contained a carboxy terminal alanine to provide more conformational flexibility. For Region 3, the sequence of ExbD residues 129-134 (K-I-G-L_132_-V-G) was chosen along with the corresponding mutant peptide representing ExbD L132Q (K-I-G-Q_132_-V-G).

Because the larger ExbD Region 2 (residues 92-121) lacks specifically identified targets and has extensive conservation of residues among most of the Gram-negative ESKAPE pathogens responsible for some of the most deadly acquired hospital infections [(Fig. 10), (67)], studies of protein-protein interactions were used to further guide the choice of residues. The sequence I-F-F-R-A-D (residues 102-107) was deemed likely to be a site of ExbD-ExbD interaction because it contains a concentration of hydrophobic residues (IFFA), which are commonly found at protein-interaction interfaces (68). In addition, three of the top five “hotspot” amino acid preferences which contribute the most binding energy in a protein interaction, were present within the semi-conserved sequence region shown in Fig. 10 [Ile, Arg, and Asp (69)]. Asp 207 was excluded from the cyclic peptide to facilitate transit through porins in the outer membrane which have a size limit of ∼ 600 da (70).

### Cyc-IFFRA inhibits TonB-dependent activity

To eliminate potential efflux of peptides out of the periplasm, strain SH03 (W3110, *tolC*::Tn*10*) was used (71). Initial rates of TonB-dependent ^55^Fe-ferrichrome transport were determined in the presence of cyclic (cyc) peptides cyc-VKVNLP and cyc-VKVNLPA from Region 1; cyc-IFFRA from Region 2; cyc-KIGLVG, and cyc-KIGQVG from Region 3 (Fig. 11). In the presence of cyc-IFFRA, the ^55^Fe-ferrichrome initial transport rate was reduced to ∼30% of the SH03 + DMSO control, indicating that it interfered with TonB system activity. Cyc-VKVNLP, cyc-VKVNLPA, cyc-KIGLVG, and cyc-KIGQVG had no effect on ^55^Fe-ferrichrome transport. These results suggested that either the peptides were not inhibitory or that they did not efficiently enter the cells. In any case the L132Q mutation results were uninterpretable since the Region 3 peptide encoding the wild-type version had no detectable effect.

**Figure 11:**
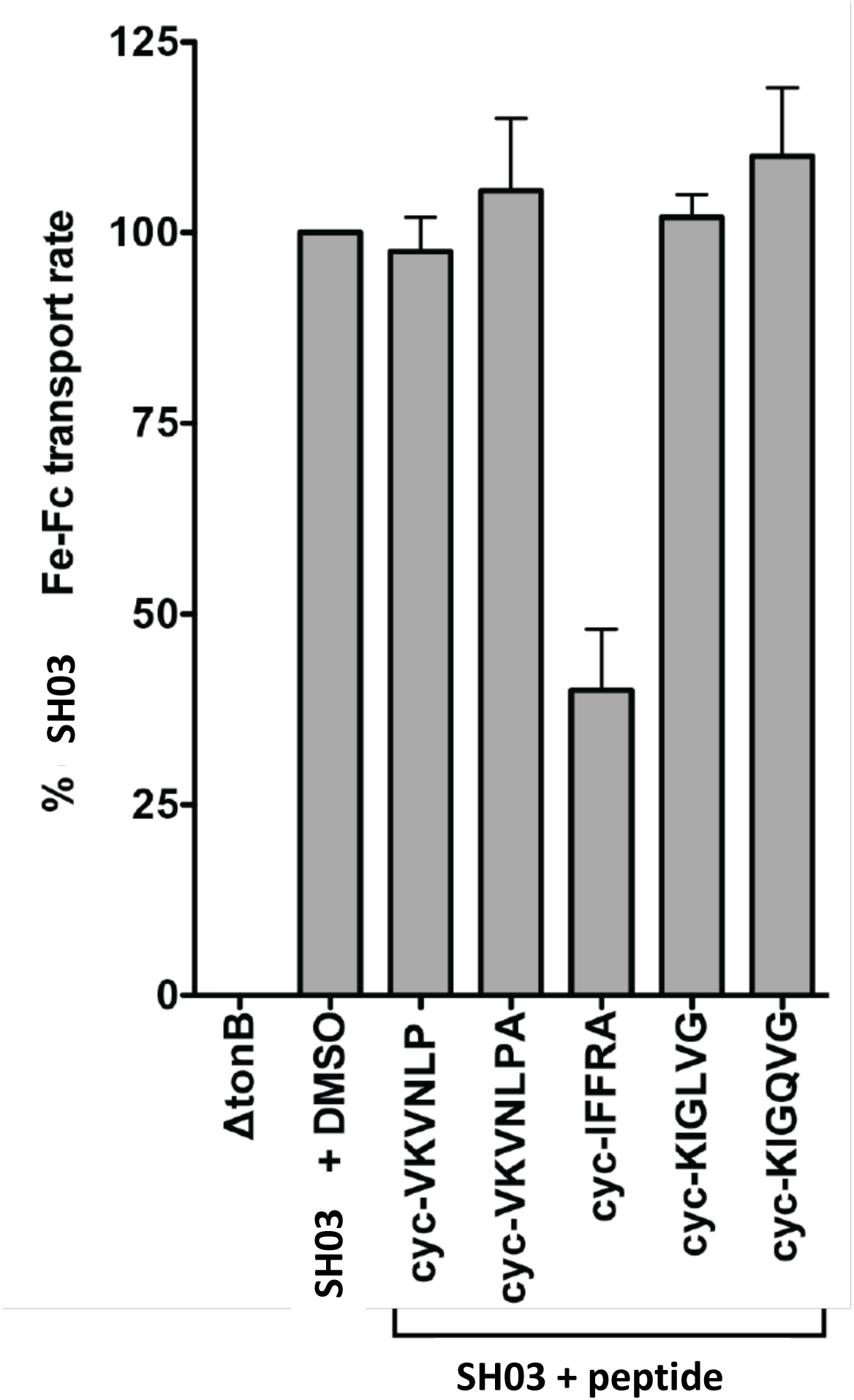
Cyclic peptide IFFRA inhibits TonB system activity. The initial rate of ^55^Fe-ferrichrome transport by strain SH03 (W3110, *tolC*::Tn*10*) in the presence of dimethyl sulfoxide (DMSO) or synthetic cyclic peptides in DMSO is shown. Cyclic peptide cyc-VKVNLP is based on ExbD residues 45-50; cyc-VKVNLPA is based on ExbD residues 45-51; cyc-IFFRA is based on ExbD residues 102-106, cyc-KIGLVG is based on ExbD residues 129-134, and cyc-KIGQVG is based on ExbD residues 129-134 with an L-to-Q substitution (see Fig. 10). Strains were grown until mid-exponential phase at which point an *A*_550_ of 1.4 cells were harvested and suspended in assay medium as previously described (115). DMSO or synthetic peptides (500 µM) suspended in DMSO were added, and samples were incubated at 30°C while shaking for five minutes prior to measuring the initial ^55^Fe-ferrichrome transport rate. The ^55^Fe-ferrichrome transport rate was recorded as the percent of the SH03 transport rate. Δ*tonB* refers to W3110 with the *tonB* gene deleted [KP1477; (112)]. The error bars indicate mean ± SEM for two biological replicates, each containing three technical replicates.

### Cyc-IFFRA inhibits formation of TonB-ExbD and, surprisingly, TonB-ExbB complexes in vivo

Stage III of the TonB energy transduction cycle is characterized by the PMF-dependent formation of formaldehyde cross-linked ExbD-TonB complexes (Fig. 2). Cys substitutions K97C and T109C on either side of the Region 2 peptide-derived sequence IFFRA capture multiple interactions with Cys substitutions in the periplasmic TonB carboxy terminus *in vivo*, and formation of those disulfides is significantly reduced or nonexistent in the PMF-unresponsive mutant, ExbD D25N (44). Taken together these results suggested that the inhibitory effect of cyc-IFFRA could be due to inhibition of Stage III of the energy transduction cycle. To test that hypothesis, strain SH03 was treated with formaldehyde in the presence of the five cyclic peptides evaluated in Fig. 11 and processed for immunoblotting with both anti-ExbD and anti-TonB antibodies (Fig. 12).

**Figure 12:**
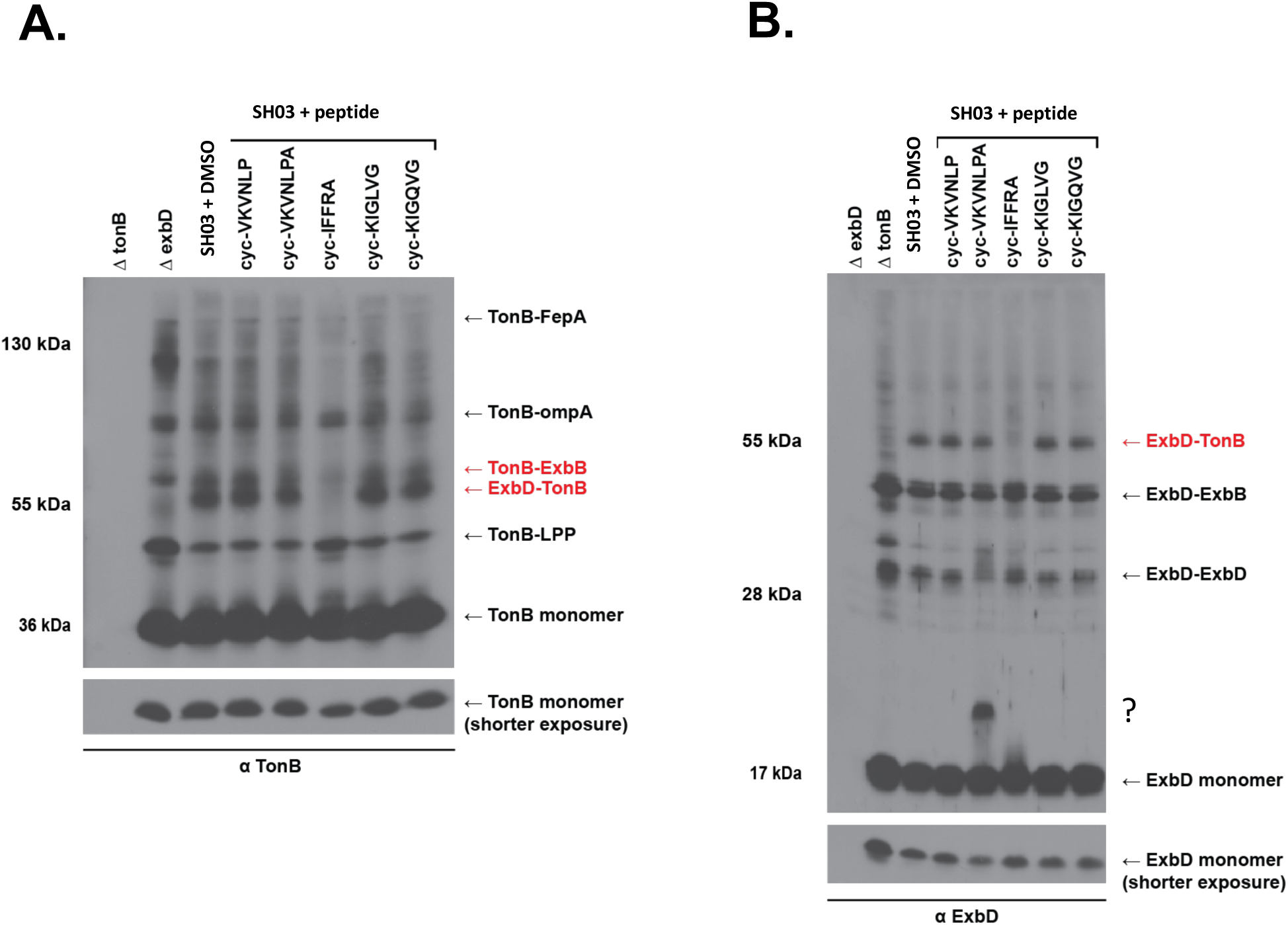
IFFRA inhibits TonB-ExbD and TonB-ExbB complex formation. Formaldehyde cross-linking of SH03 (W3110, *tolC*::Tn*10*) in the presence of dimethyl sulfoxide (DMSO) or synthetic cyclic peptides in DMSO is shown. Cyclic peptide cyc-VKVNLP is based on ExbD residues 45-50; cyc-VKVNLPA is based on ExbD residues 45-51; cyc-IFFRA is based on ExbD residues 102-106, cyc-KIGLVG is based on ExbD residues 129-134, and cyc-KIGQVG is based on ExbD residues 129-134 with an L-to-Q substitution (see Fig. 10). Strains were grown to mid-exponential phase, at which point cells were harvested and suspended in 100 mM sodium phosphate buffer (pH 6.8). DMSO or the synthetic cyclic peptides (500 µM in DMSO) were added five minutes prior to formaldehyde cross-linking as previously described (122). Proteins from equivalent numbers of cells from the same cultures were analyzed on immunoblots of SDS 13% polyacrylamide gels with (A) anti-TonB monoclonal antibodies and (B) anti-ExbD polyclonal antibodies. Identity of samples are indicated at the top of both immunoblots. Δ*exbD* refers to KP1045; Δ*tonB* refers to KP1477. (**A.**) Positions of the previously characterized TonB formaldehyde cross-linked complexes are indicated on the right: TonB-FepA, TonB-OmpA, TonB-ExbB, PMF-dependent TonB-ExbD, and the TonB-LPP complex (14, 28, 29). The position of TonB monomer is also indicated. Mass markers are indicated on the left. (**B.**) Positions of the previously characterized ExbD formaldehyde cross-linked complexes are indicated on the right: PMF-dependent ExbD-TonB complex, ExbD-ExbB complex, and ExbD-ExbD homodimer (29). The position of ExbD monomer is also indicated. The position of a potential complex between monomeric ExbD and the cyc-VKVNLPA peptide is indicated by “?”. Mass markers are indicated on the left. Red text identifies the position of TonB-ExbB and ExbD-TonB complexes that are inhibited in the presence of cyc-IFFRA. Beneath A and B are shorter exposures of the immunoblots showing relative levels of monomers.

Consistent with its inhibition of iron transport in Fig. 11, the Region 2 cyc-IFFRA peptide prevented the Stage III TonB-ExbD complex detected with both anti-TonB antibody (Fig. 12A) and anti-ExbD antibody (Fig. 12B). It also unexpectedly decreased formation of TonB-ExbB complexes compared to the control (SH03 + DMSO), but not TonB-Lpp, TonB-OmpA or TonB-FepA complexes. Formaldehyde cross-linking of TonB with Lpp and OmpA appear to be off-pathway contacts made by unenergized TonB since a double *lpp, ompA* mutant has only slightly reduced iron transport (14). The FepA complex likewise represents an inactive TonB-FepA complex which occurs with unenergized TonB (72). The cyc-IFFRA peptide had no effect on ExbD homodimer or ExbD-ExbB heterodimer formation.

Inhibition of the TonB-ExbB complex by a cyclic peptide corresponding to a distal periplasmic region of ExbD suggested that this region of ExbD played a role in assembly or conformation of the TonB-ExbB interaction *in vivo*. While monomeric formaldehyde is a probe for close contacts (73), the absence of a formaldehyde-cross-linked complex only means that the formaldehyde-cross-linkable residues have moved out of correct apposition--due either to dissolution of the complex or to a change in conformation of one or more proteins within the complex.

The cyc-VKVNLPA peptide from Region 1 appeared to formaldehyde cross-link to ExbD monomer (labeled “?”) and disrupt ExbD homodimer cross-linking (Fig. 12B), consistent with our identification of ExbD homodimerization through Region 1 residues in full-length ExbD (59). Monomeric formaldehyde inserts a CH_2_ group between two reactive residues (K, Y, H, C ⋙ R, W) (74). It seems likely that cyc-VKVNLPA formaldehyde cross-linked to ExbD monomer through its Lys residue. None of the other peptides contained strongly cross-linkable residues, which explains why binding to ExbD monomer was not detected with the inhibitory cyc-IFFRA peptide from Region 2. If cyc-VKVNLPA did cross-link to ExbD monomer, it means that it was able to penetrate the outer membrane, even though it was the largest of the peptides. This result suggested how sensitive translocation across the outer membrane, presumably via porin proteins, was to overall conformation because the smaller version of the same peptide that also contained a cross-linkable Lys residue apparently did not enter the periplasm. Alternatively, it did, but could not bind to ExbD. Because cyc-VKVNLPA did not affect the iron transport rate in Fig. 11, it was not a candidate to be a TonB system inhibitor. Cyc-IFFRA, however, showed promise.

## DISCUSSION

Outer membrane active transporters of Gram-negative bacteria, including ESKAPE pathogens, are powered by the TonB-system (75), which is one of three systems in *E. coli* K12 that each use two similar cytoplasmic membrane proteins and the PMF to power cell envelope events: ExbB and ExbD of the TonB system energize active transport across the outer membrane (76); TolQ and TolR power proper outer membrane invagination (77); MotA and MotB act as stators for flagellar rotation (78). Of the three systems, the TonB system is the least complex so far with ExbB/D and TonB located in the cytoplasmic membrane and a variety of TBDTs in the outer membrane (Fig. 1), although there is evidence for additional mystery proteins (79, 80). What they all have in common are residue similarities in the last two transmembrane domains of ExbB/TolQ/MotA and an invariant Asp residue in approximately the same position in the single transmembrane domain of ExbD/TolR/MotB. Other than similarities among their transmembrane domains, the remaining residues most likely account for their disparate functions. The overall mechanism of TonB-mediated energy transduction remains a mystery. Results from the also-mysterious Mot and Tol systems provide informative comparisons.

*In vivo* studies of the paralogue MotB transmembrane domain demonstrate the importance of Asp32 (D25 of ExbD; D23 of TolR) to flagellar rotation, where out of 15 substitutions only the conservative replacement D32E shows a slight amount of activity (49). MotB (308 residues) forms homodimers through a face on its single α-helical transmembrane domain (46, 81). MotB also binds to peptidoglycan through residues ∼ 209-227, a logical function for a flagellar rotation stator (82, 83). The need for a “plug” (MotB residues 52-65, C-terminal to the transmembrane domain) is explained if unassembled MotA/B complexes leak protons until they are assembled in the flagellar motor (84).

*In vivo* studies on paralogue TolR (142 residues in length) show that two TolR transmembrane domains homodimerize in a pattern similar to the MotB transmembrane domains (Fig. 3B). Like MotB, TolR has a conserved Asp residue in its transmembrane domain that is inactivated by the D23R substitution, while the conservative D23E substitution retains lower but detectable activity (48). ExbB/D and TolQ/R are the most closely related paralogues and can in fact substitute (∼ 10%) for one another in their respective interactions with either non-cognate TonB or Tol-Pal proteins (45, 53–55, 85). Cys substitution Y117C in the periplasmic carboxy terminus traps TolR such that it cannot undergo needed conformational transformations (86). Similarly, disulfide-linked homodimers through the corresponding residue in ExbD (Y112C) are inactive (36). Both results speak to the dynamic nature of their respective periplasmic domain interactions.

Unlike the MotB transmembrane domains that apparently retain a static relationship and thereby support binding of its periplasmic domain to peptidoglycan, homodimeric TolR transmembrane domains appear to rotate relative to one another. Cys substitutions on opposite sides of the transmembrane domain helix [L22 and V24 (corresponding to ExbD I24 and V26, Fig. 3B),] trap TolR as disulfide-linked homodimers that inactivate the Tol system. Addition of the reducing agent DTT reduces the disulfide bond and restores activity (47). This requirement for movement raises a question whether the TolR periplasmic domain binds to peptidoglycan *in vivo*. Consistent with that, TolR is less than half the length of MotB, seemingly making it less likely to access the peptidoglycan. In addition, TolR also does not have an identifiable plug. In a contrasting view, TolR is hypothesized to reach the peptidoglycan as a disordered protein (87). Cryptic peptidoglycan binding is observed for a truncated TolR periplasmic domain, but it is not clear how the binding affinity compares to that of canonical OmpA peptidoglycan binding (88).

### The ExbD transmembrane domain

At 141 residues, ExbD is essentially the same length as TolR (45). Braun et al. were the first to show that ExbD D25N is inactive (43). Our *in vivo* studies described here on the predicted ExbD transmembrane domain (residues 23-43) were late to the party but were somewhat more thorough from a mutagenic point of view. They first of all indicated that, like the MotB and TolR transmembrane domains, the ExbD transmembrane domain homdimerized in an α-helical configuration.

All residues in the ExbD transmembrane domain except Asp25 could be substituted with Cys and retain function in colicin and phage sensitivity assays. Replacement by none of the remaining 19 amino acids at residue 25 except the conservative replacement D25E supported any activity, thus aligning well with the importance of the transmembrane domain Asp in MotB and TolR (48, 49). For the first time, alternative positions for Asp25 within the transmembrane domain were assessed, showing that ExbD Asp25 did not function at any other position within the transmembrane domain. Because participation of the TonB and ExbB transmembrane domains in PMF-responsiveness is ruled out, it appears that in the TonB system PMF is engaged solely through ExbD Asp25 (26, 27). Consistent with that, progression to Stage III in the energy transduction cycle is prevented either by the protonophore CCCP (carbonyl cyanide *m*-chlorophenyl hydrazone) that collapses PMF or by the unresponsive *exbD* D25N mutation [Fig. 2; (29, 42)].

### Two different domains in the ExbD amino terminal region

Residues 23-43 were originally proposed to constitute the ExbD transmembrane domain (20). Our results suggested instead that the functional boundaries of the ExbD transmembrane domain could be revised to extend only from ExbD F23 to P39. A set of three ExbD Cys substitutions in this region appeared to represent an α-helical configuration, requiring both oxidant and the presence of ExbB to form disulfide-linked homodimers through those residues. This revision somewhat positions the ExbD transmembrane domain between two prolines at residues 22 and 39, that are conserved in TolR but not MotB (Fig. 3).

In contrast, each sequential residue from L40C through V43C homodimerized strongly without the requirement for either added oxidant or ExbB, suggesting that they constituted an intrinsically disordered domain. [ExbD T42C did not homodimerize in the absence of ExbB because it appeared to be chemically modified.] Thus, ExbD residues L40-V43 appeared to initiate the previously identified intrinsically disordered region of ExbD that terminates with residue 63, homodimerizes, and contains the conserved motif V45-V47-L49-P50 that is required for signal transduction of PMF to the TonB periplasmic domain (59, 60).

The ExbD_23-63_-TOXCAT fusion protein, where none of the residues was Cys-substituted, homodimerized either through ExbD transmembrane domain residues (residues 23-39) or an apparently disordered region (residues 40-63), or both. That result ruled out a requirement for participation of the Dsb system in ExbD_23-63_ homodimerization. The ExbD_23-63_-TOXCAT fusion protein homodimerization did not require ExbB, indicating an intrinsic ability to homodimerize that was consistent with previous studies identifying homodimerization through residues 44-56 using *in vivo* photo-cross-linking (59) and disulfide-linked homodimerization of ExbD through Cys substitutions in its periplasmic domain (residues 92-121) that also does not require ExbB *in vivo* (36). It thus appeared that both disordered and ordered regions of the ExbD periplasmic domain homodimerized independently of ExbB and it is likely that the observed homodimerization of ExbD_23-63_-TOXCAT both in the presence and absence of ExbB was due to the presence of the disordered region and not the transmembrane domain. If the entire periplasmic domain of ExbD homodimerizes independently of ExbB, perhaps it might initiate formation of ExbD transmembrane domain homodimers rather than vice versa.

### Required motion of the ExbD transmembrane domains

Because L40C-V43C homodimerized in the absence of ExbB, it suggested that the ExbD residues 23C-39C were likely also homodimerized due to their proximity in the absence of ExbB, but without formation of disulfide-linked homodimers. Combined, these results suggested that the ExbD transmembrane domains had the capacity to homodimerize in two different configurations *in vivo* which could be consistent with a requirement for motion. The importance of motion was confirmed when spontaneously disulfide-linked Cys substitutions at positions T42 and V43 inactivated TonB-dependent transport and reduction of the disulfides reactivated it. These results mirrored similar approaches showing that TolR transmembrane domains rotate relative to one another (47).

Results from *in vivo* photo-cross-linking studies suggested that the motion could be rotation because sites on all faces of the ExbD transmembrane domain participated in formation of homodimers. Consistent with that idea, *in vivo* photo-cross-linking also showed that all sites investigated in the ExbD transmembrane domain formed photo-cross-linked complexes with ExbB.

Thus, two different techniques have now identified the likely rotation of ExbD/TolR transmembrane domain homodimers relative to one another *in vivo*. By assessing disulfide-linked homodimers, Celia et al. also suggested that the ExbD transmembrane helices rotated with respect to one another (89).

### Results in the context of recent subcomplex cryo-EM structures

The recent cryo-EM structures of ExbB/D and MotA/B from several species show them to be asymmetric subcomplexes of 5 ExbB/MotA and 2 ExbD/MotB (51, 52, 89). The congruence of the proposed structures is striking, especially since it is seen across different species and with the GldL/M proteins involved in gliding motility of *Flavobacterium johnsoniae* (90). Authors of the MotA/B cryo-EM papers suggest that MotA rotates around MotB homodimer, held static by interaction with peptidoglycan (51, 52). Hennell-James et al. suggests that the GldL pentamer rotates relative to the GldM homodimer (like MotB) (90). It has been suggested that these admittedly beautiful structures provide the key to the mechanism of the protein complexes that all use PMF for energy (91) and that ExbB/D follows the same mechanism as MotA/B (51).

An argument can be made for the validity of the MotAB structure because the first transmembrane domain of MotA, which has four rather than three transmembrane domains like ExbB and TolQ, stands in for their respective TonB/TolA transmembrane domains, thus obviating concerns that TonB is not part of the ExbB/D cryo-EM structures and providing reassurance for the other structures that, because they all exhibit the same 5:2 asymmetric architecture, they accurately reflect the *in vivo* state of the complexes. Nonetheless, there is still a need to understand how and where TonB/TolA fit into ExbB,TolQ/ExbD,TolR complexes and to address the cellular ratios of 7 ExbB/TolQ:4 ExbD/TolR:1TonB/TolA (11, 48, 92).

More remains to be discovered about the mechanism of these proteins. As Deme et al. states “No ion permeation across the bilayer was seen in any simulation, supporting the idea that the complexes currently seen will require rearrangement for activity.” (51). Celia et al. likewise did not see a channel in ExbB5/D2 through which a proton could be translocated and suggest that structural plasticity is likely important (89). In support of a general need to incorporate *in vivo* results into interpretations based on structural information, recent data from Nilaweera et al. demonstrate the importance of studying the TonB system and its outer membrane TBDTs in their native cellular environment compared to *in vitro* studies (93).

Evidence for the *in vivo* existence of ExbB tetramers and MotA tetramers (as opposed to pentamers) is equally persuasive, and has the advantage that all required proteins, known and unknown, are present in whole cells (79–81). For example, ExbB formaldehyde cross-links as a dimer of homodimers *in vivo* (80). In another example, ExbD photo-cross-links to TonB *in vivo* through residues that boundary its transmembrane domain suggesting that the ExbD and TonB transmembrane domains are at some point adjacent (63), a result seemingly at odds with the structural data where the lumen of an ExbB pentamer cannot accommodate additional transmembrane domains beyond the two of ExbD. Heterodimerization of TonB-ExbD transmembrane domains would be consistent with Stage II in the proposed energy transduction cycle (Fig. 2).

While *in vivo* studies of the TonB system have admittedly not always been interpreted correctly [see (39) subsequently corrected in (94)], it is also worth noting that structural studies have failed to correctly represent *in vivo* conformations of TonB carboxy terminal homodimers (41) and important interactions between the TonB carboxy terminus and TBDTs FhuA and BtuB proteins (40). Structural studies have also missed interactions between the TonB carboxy terminal amphipathic helix and TBDT FepA (18). Additionally, Cys substitutions that formed ExbD disulfide-linked homodimers cannot be modeled on the ExbD NMR structure without significant distortion (36). It seems possible that elements of the model based on *in vivo* data summarized in Fig. 2 could provide important insights that have not otherwise been gained by structural analysis. Also, considering the parable of the blind men and the elephant, could there be *in vivo* equilibria among the several proposed ratios of TonB:ExbB:ExbD (11, 25, 80, 95, 96)?

Given the complexity and dynamic nature of the differing protein-protein interactions that occur in each system, a true understanding of their mechanisms awaits *in vivo* data and *in vitro* structural models that are significantly more congruent. Currently, much of the *in vivo* data for all three PMF-dependent systems remains to be rationally incorporated into models arising from structural analyses.

### ExbD L132 is a non-essential residue

Braun et al. isolated a spontaneous ExbD L132Q mutant that was inactive, the first to demonstrate the importance of ExbD in all TonB system activities (43). In this work, the construction and expression of all possible amino acid substitutions at *exbD* codon 132 showed that L132 is not a mechanistically important residue, even though L132Q prevents formaldehyde crosslinking to TonB and stalls the energy transduction cycle at Stage I [Fig. 2; (42)]. Because other structurally diverse substitutions such as L132C and L132W were fully active, the L132Q mutation likely caused structural distortion that prevented productive ExbD-TonB interaction.

### ExbD as a target for a novel antibiotic

The TonB system has been recognized as a potential target for novel antibiotics for many years because bacteria need sufficient iron to mount infection in their hosts, which have evolved to sequester it from them (97–103). There appears to be an escalating battle between bacterial pathogen and host, starting with the deliberately low levels of iron in body fluids that can be thwarted by the specialized high-affinity iron acquisition systems such as the TonB system that bacteria use to retrieve it (9).

TonB protein undergoes three different protein-protein interactions during an energy transduction cycle: TonB-TonB, TonB-ExbD, and TonB-TBDT (Fig. 2). Clues that these could constitute targets for intervention arise because overexpression of TonB exhibits a dominant negative gene dosage effect. Proteolytic fragments of TonB that are produced when the level of TonB exceeds the level of ExbB needed to proteolytically stabilize it can competitively inhibit energy transduction (104, 105). An exogenously added peptide derived from a conserved consensus sequence known as the “TonB box” in TBDTs showed promise (106) but was not examined further.

Shotgun and rational design approaches have also been used to search for compounds that could inhibit the TonB system. High-throughput screening of small molecule libraries have identified compounds that inhibit TonB system activity (107, 108). However, none of the compounds was shown to directly target the TonB system and potential modes of inhibition were not identified.

Compounds that inhibit individual TonB-TBDT interactions might not be therapeutically optimal since TonB interacts differently with the several different TBDTs of *E. coli* K12, which could potentially diminish or eliminate the inhibition (41, 109). The essential TonB-ExbD interaction appeared to be a better target since each protein and their interaction are required for energy transduction through all TBDTs (Fig. 2).

We were expecting to discover that a peptide that is inhibitory when supplied endogenously [residues 44-63; (63)] would also be inhibitory if supplied exogenously as a cyclic peptide (residues 45-51), but that was not the case. Possibly it had no effect because it was too short or possibly because an insufficient amount was able to cross the outer membrane and reach the periplasm. Instead, the cyclic peptide cyc-IFFRA, derived from ExbD periplasmic domain residues 102-106 in a region that interacted with the periplasmic domain of TonB prevented the formation of the essential ExbD-TonB complex *in vivo* and inhibited TonB system activity. Because the peptide was cyclic and lacked an OmpT recognition site, it appeared to have largely survived proteolysis by envelope proteases. It would be interesting to survey the effect of this peptide on ESKAPE or other Gram-negative pathogens. The use of customized cyclic peptides to inhibit the essential ExbD-TonB interaction in Gram-negative bacteria could serve as a prototype for future antibiotic development.

## MATERIALS AND METHODS

### Strains and plasmids

Plasmids used in this work are listed in Table 5. Transmembrane domain codons 23-43 of *exbD* were engineered individually to Cys codons by site-directed mutagenesis PCR on plasmid pKP999 which encodes wild type ExbD expressed from a propionate-inducible promoter (29). Open reading frames of all *exbD* mutants were sequenced at the Penn State Genomics Core Facility, University Park, PA to rule out unintended base changes. The various individual Cys-substituted mutants were transformed to strains RA1045 [(*11exbD, 11tolQR*); (54)], RA1017 [(*11exbBD, 11tolQRA*); (110)], and KP1509 [(*11exbD, 11tolQR, ΔtonB*); (29)] as described in the figure legends. All three strains are derivatives of W3110 (111). For studies of ExbD cyclic peptides, strain SH03 (W3110, *tolC*::Tn*10*) and KP1477 [W3110, Δ*tonB* (112)] were used.

**Table 5.** Plasmids used in this study

| **Plasmid** | **Relevant phenotype** | **Source or reference** |
| --- | --- | --- |
| pccKan | TOXCAT system vector | (58) |
| pccGlpA | GlpA-TOXCAT fusion | (58) |
| pccGlpAG83I | GlpA G83I-TOXCAT fusion | (58) |
| pKP523 | ExbD<sub>23-63</sub>-TOXCAT fusion | (58) |
| pKP568 | TonB(C18G) | (72) |
| pKP945 | TonB(C18G, A150C) | (29) |
| pKP999 | pPro24 expressing wild-type ExbD | (29) |
| pKP1000 | ExbD(A92C) | (29) |
| pKP1064 | ExbD(D25N) | (29) |
| pKP1394 | ExbD(F23C) | Present study |
| pKP1264 | ExbD(I24C) | Present study |
| pKP1265 | ExbD(D25C) | Present study |
| pKP1284 | ExbD(V26C) | Present study |
| pKP1285 | ExbD(M27C) | Present study |
| pKP1286 | ExbD(L28C) | Present study |
| pKP1395 | ExbD(V29C) | Present study |
| pKP1396 | ExbD(L30C) | Present study |
| pKP1405 | ExbD(L31C) | Present study |
| pKP1397 | ExbD(I32C) | Present study |
| pKP1406 | ExbD(I33C) | Present study |
| pKP1389 | ExbD(F34C) | Present study |
| pKP1386 | ExbD(M35C) | Present study |
| pKP1399 | ExbD(V36C) | Present study |
| pKP1388 | ExbD(A37C) | Present study |
| pKP1390 | ExbD(A38C) | Present study |
| pKP1387 | ExbD(P39C) | Present study |
| pKP1398 | ExbD(L40C) | Present study |
| pKP1391 | ExbD(A41C) | Present study |
| pKP1392 | ExbD(T42C) | Present study |
| pKP1393 | ExbD(V43C) | Present study |
| pKP1412 | ExbD(D25N, L40C) | Present study |
| pKP1413 | ExbD(D25N, A41C) | Present study |
| pKP1445 | ExbD(D25N, T42C) | Present study |
| pKP1406 | ExbD(D25N, V43C) | Present study |
| pKP1191 | ExbD(D25A) | Present study |
| pKP1276 | ExbD(D25A, F23D) | Present study |
| pKP1273 | ExbD(D25A, I24D) | Present study |
| pKP1201 | ExbD(D25A, V26D) | Present study |
| pKP1213 | ExbD(D25A, M27D) | Present study |
| pKP1229 | ExbD(D25A, L28D) | Present study |
| pKP1202 | ExbD(D25A, V29D) | Present study |
| pKP1413 | ExbD(D25A, L30D) | Present study |
| pKP1272 | ExbD(D25A, L31D) | Present study |
| pKP1277 | ExbD(D25A, I32D) | Present study |
| pKP1287 | ExbD(D25A, I33D) | Present study |
| pKP1288 | ExbD(D25A, F34D) | Present study |
| pKP1322 | ExbD(D25A, M35D) | Present study |
| pKP1351 | ExbD(D25A, V36D) | Present study |
| pKP1359 | ExbD(D25A, A37D) | Present study |
| pKP1349 | ExbD(D25A, A38D) | Present study |
| pKP1360 | ExbD(D25A, P39D) | Present study |
| pKP1214 | ExbD(D25A, L40D) | Present study |
| pKP1215 | ExbD(D25A, A41D) | Present study |
| pKP1199 | ExbD(D25A, T42D) | Present study |
| pKP1197 | ExbD(D25A, V43D) | Present study |
| pKP2234 | ExbD(T21am) | Present study |
| pKP2235 | ExbD(F23am) | Present study |
| pKP2064 | ExbD(I24am) | Present study |
| pKP2236 | ExbD(D25am) | Present study |
| pKP2229 | ExbD(M27am) | Present study |
| pKP2224 | ExbD(V29am) | Present study |
| pKP2237 | ExbD(I32am) | Present study |
| pKP2065 | ExbD(F34am) | Present study |
| pKP2225 | ExbD(M35am) | Present study |
| pKP2226 | ExbD(A37am) | Present study |

### Media and culture conditions

Luria-Bertani (LB), tryptone (T), and M9 minimal salts media were prepared as described previously (113, 114). Except for plasmid-less strains, all liquid cultures, agar plates, and T-top agar were supplemented with 35 *µ*g of chloramphenicol/ml and plasmid-specific levels of propionate were added during the assay for the expression of ExbD cysteine substituted proteins at chromosomal levels as determined by immunoblot. M9 salts were supplemented with 1.0% glycerol (wt/vol), 0.4 mg of thiamine/ml, 1 mM MgSO_4_, 0.5 mM CaCl_2_, 0.2% Casamino Acids (wt/vol), 40 mg of tryptophan/ml, and 1.85 *µ*M FeCl_3_. Cultures were grown at 37°C with continuous aeration.

### Sensitivity to colicins and bacteriophage

To measure the relative activity of ExbD mutants, strains expressing ExbD mutants at chromosomal levels were grown to mid-logarithmic phase in T-broth and plated in T-top agar on T-agar plates containing 35 *µ*g of chloramphenicol/ml. The appropriate concentration of propionate inducer required for chromosomal level expression of each mutant was determined by immunoblotting and was present in growth media throughout the assay. 5-fold and 10-fold dilutions of colicin or bacteriophage, respectively, were spotted on the inoculated T-top layer. The zones of clearing were recorded after 18 h incubation at 37°C (56, 115).

### Iodine mediated and spontaneous disulfide cross-linking

Cells grown overnight in LB were subcultured 1:100 into T-broth with various amounts of Na-propionate to achieve chromosomal levels of expression (from 0.0-1.0 mM and in the case of ExbD V29C, glucose at 0.005% was added to repress expression). At mid-exponential phase, 0.8 A_550_-ml of cells were pelleted, washed and suspended in 100 *µ*l of cross-linking buffer (20mM sodium phosphate, pH 7.4, 150mM NaCl). Then cells were treated with 0.2 mM iodine (I_2_) from a 20 mM stock solution in ethanol and kept on ice for 10 min. Unreacted free Cys residues were blocked with 20 mM *N*-ethylmaleimide (NEM) for 5 min on ice (46). Cells were then precipitated on ice with equal volume of 20% TCA. Pellets were suspended in SDS gel sample buffer supplemented with 50mM iodoacetamide and without β-mercaptoethanol (β-ME) (116). To check the level of protein expression, 0.2 A_550_-ml of mid-exponential phase cells from the experimental samples were harvested immediately on ice with equal volumes of 20% TCA and solubilized in SDS gel sample buffer containing β-ME. A kinetic study of dimer formation showed completion of iodine-mediated disulfide cross-linking within 15 sec of incubation on ice (data not shown). Both iodine-treated non-reduced and β-ME-treated reduced samples were resolved on 15% SDS-polyacrylamide gels and immunoblotted with ExbD-specific polyclonal antibody (11). Equal loads were confirmed by Coomassie staining of the PVDF membrane used for the immunoblot. For spontaneous disulfide cross-linking, iodine and NEM were omitted.

### In vivo photo-cross-linking

Plasmid pEVOL encodes an orthogonal tRNA synthetase and a corresponding orthogonal tRNACUA that recognizes amber (UAG) codons (117). Plasmids with amber substitutions in *exbD* were engineered, co-expressed with pEVOL in RA1045 (W3110, Δ*exbD*, Δ*tolQR*) and processed as previously described (17). At A_550_ of ∼0.2, sub-cultures in supplemented M9 minimal media were treated with 0.004**%** arabinose to induce expression of the tRNA synthetase/tRNA pair from pEVOL, 0.8 mM of the photo-cross-linkable amino acid p-Benzoyl-L-phenylalanine (pBpa; Bachem AG, www.bachem.com) and levels of propionate required for ExbD protein expression at near-chromosomal levels. 2.0 A_550_-ml aliquots were harvested at A_550_ ∼ 0.5, pelleted at room temperature and suspended in 2.0 ml supplemented M9 media (as described above with 1.85 μM FeCl_3_ instead of 37 μM, and without antibiotics, pBpa, and propionate). Equal volumes of each sample were transferred into single wells of each of two 24-well plates (NEST Cell Culture Plate, cat: 702001). One plate was irradiated with 365-nm light for 30 min on ice and the second plate was simultaneously incubated in the dark for 30 min on ice. Samples were precipitated with an equal volume of 20% TCA as described previously and processed for immunoblot analysis with ExbD-specific polyclonal antibodies and TonB-specific monoclonal antibodies (11, 118). Equal loads were confirmed by Coomassie staining of the PVDF membrane used for the immunoblot.

### Initial rates of [^55^Fe]-ferrichrome transport

The initial rates of iron transport were determined as described previously (56, 115). Mid-exponential-phase subcultures were pelleted and suspended in iron transport assay media, and the initial rates of [^55^Fe]-ferrichrome transport were determined. Equal proportions of the suspended cells were precipitated with an equal volume of 20% TCA to confirm the chromosomal level expression of ExbD mutants by immunoblot. Precipitates were solubilized in SDS gel sample buffer (116), electrophoresed on a 15% SDS polyacrylamide gel and immunobotted with anti-ExbD polyclonal antibody (11). Inhibitory ExbD-based cyclic peptides were purchased from GenScript USA, Inc., Piscataway, New Jersey. Assay of equal numbers of cells in each sample was confirmed by Coomassie staining of the PVDF membrane used for the immunoblots.

### In-vivo formaldehyde cross-linking

Cultures were grown overnight to saturation in LB broth at 37°C with aeration. The saturated cultures were subsequently subcultured 1:100 in supplemented M9 minimal media. At mid-exponential phase, volumes were adjusted to harvest 0.5 A_550_-ml of cells. Bacteria were suspended in sodium phosphate buffer (pH 6.8) and treated for 15 min at room temperature with 1% monomeric formaldehyde (Electron Microscopy Sciences Cat #15710) as previously described (29). Following formaldehyde treatment, bacteria were pelleted and suspended in SDS gel sample buffer (116), with equal cell numbers loaded on SDS 13% polyacrylamide gels. ExbD complexes and TonB complexes were detected by immunoblotting with ExbD-specific polyclonal antibodies and TonB-specific monoclonal antibodies, respectively (11, 118). Equal loads were confirmed by Coomassie staining of the PVDF membrane used for the immunoblot.

## Acknowledgements

We thank Ann Ollis and Christine Dubowy for construction and assay of the Cys relocation mutants. We thank Yan Shipelskiy for construction and assay of ExbD D25X mutants. We thank Garrick Treaster, Anne Ollis, and Elise Zhao for construction and characterization of mutations at ExbD residue L132. We thank Erica Johnson for construction of several amber mutations in *exbD*. We thank William Russ and Donald Engelman for TOXCAT vectors and *E. coli* NT326 cells, and Kevin Bertrand and Stuart Huntley for the gift of SH03. We thank Ray Larsen for editorial feedback on the manuscript. Support from NIAID R21 AI113622, NIGMS R01 GM112710, and Violet S. Postle is gratefully acknowledged.

